# Single-nucleus RNA/ATAC-seq in early-stage HCM models predicts SWI/SNF-activation in mutant-myocytes, and allele-specific differences in fibroblasts

**DOI:** 10.1101/2024.04.24.589078

**Authors:** Tilo Thottakara, Arun Padmanabhan, Talha Tanriverdi, Tharika Thambidurai, Jose A. Diaz-RG, Sanika R. Amonkar, Jeffrey E. Olgin, Carlin S. Long, M. Roselle Abraham

## Abstract

Hypertrophic cardiomyopathy (HCM) is associated with phenotypic variability. To gain insights into transcriptional regulation of cardiac phenotype, single-nucleus linked RNA-/ATAC-seq was performed in 5-week-old control mouse-hearts (WT) and two HCM-models (R92W-TnT, R403Q-MyHC) that exhibit differences in heart size/function and fibrosis; mutant data was compared to WT. Analysis of 23,304 nuclei from mutant hearts, and 17,669 nuclei from WT, revealed similar dysregulation of gene expression, activation of AP-1 TFs (FOS, JUN) and the SWI/SNF complex in both mutant ventricular-myocytes. In contrast, marked differences were observed between mutants, for gene expression/TF enrichment, in fibroblasts, macrophages, endothelial cells. Cellchat predicted activation of pro-hypertrophic IGF-signaling in both mutant ventricular-myocytes, and profibrotic TGFβ-signaling only in mutant-TnT fibroblasts. In summary, our bioinformatics analyses suggest that activation of IGF-signaling, AP-1 TFs and the SWI/SNF chromatin remodeler complex promotes myocyte hypertrophy in early-stage HCM. Selective activation of TGFβ-signaling in mutant-TnT fibroblasts contributes to genotype-specific differences in cardiac fibrosis.

## INTRODUCTION

Hypertrophic cardiomyopathy (HCM) is the most common inherited cardiomyopathy^1^, and cause of sudden cardiac death in young individuals. A large proportion of HCM patients have ≥1 pathogenic variants in sarcomeric protein genes, that promote myocyte hypertrophy, cardiac fibrosis, coronary microvascular remodeling^2^, and lead to left ventricular hypertrophy (LVH), LV hypercontractility, diastolic dysfunction, microvascular ischemia^3^, left atrial enlargement^4^. Notably, the location and degree of cardiac hypertrophy, fibrosis, as well as the age of onset of symptoms such as dyspnea, angina, heart failure, arrhythmias, is highly variable^5^. But the transcriptional regulation of phenotypic heterogeneity in HCM is poorly understood.

In order to understand molecular pathophysiology in HCM, several profiling studies (single nucleus RNA-seq^6, 7^, bulk RNA-seq^8–10^, proteomics, metabolomics^11–13^) have been performed in heart tissue from genotyped HCM patients at established disease stage, undergoing septal myectomy or heart transplant. These studies reveal marked changes in the transcriptome, metabolome and proteome, that are independent of HCM genotype. This is surprising because pathogenic variants in sarcomeric protein genes often have distinct biophysical effects^14^ that would be expected to lead to transcriptional differences. It is unknown whether the transcriptional landscape in HCM is similar at early and late disease stages for each pathogenic variant and clinical phenotype, or whether genotype-specific activation of transcriptional programs evolves into a common molecular phenotype, due to transcriptional convergence^6^, at established disease stage. Insights into transcriptional regulation of phenotypic heterogeneity are needed to assist with development of precision therapies for prevention of cardiac remodeling and complications such as heart failure and arrhythmias.

Single-nucleus (sn)-RNA-seq and ATAC-seq are powerful tools to study transcriptional changes and their regulation at a single cell level in all cell types in the heart, without cell sorting or dissociation (which can lead to biases associated with cardiomyocyte fragility). In this study, we employed the commercially available Multiome Kit to obtain linked gene expression and chromatin accessibility data from the same nuclei in two extensively characterized HCM mouse models^9, 10, 15^ that express pathogenic variants in thin (*Tnnt2)*^16, 17^ or thick filament (*Myh6*)^18^ genes, leading to phenotypes that span the spectrum of human disease (mild/no clinical LVH, sudden death at a young age^19^ in R92W-TnT^+/-^, and moderate LVH with heart failure requiring transplantation in middle age^20, 21^ in R403-βMyHC^+/-^). Both mutant proteins are incorporated into sarcomeres^16, 22^ and increase tension cost, but have markedly different biophysical effects^14^. Residue-92 is located in the α-tropomyosin-binding domain of cardiac troponin T, whereas residue-403 is located in the actin-binding motor domain of myosin heavy chain^23^. R92W-TnT^+/-^ confers increased myofilament Ca^2+^ sensitivity of force development^16^, and is associated with LV hypercontractility, diastolic dysfunction, lower LV mass^16^, interstitial fibrosis, and induction of the fetal gene program^16^ in mice. Studies in R403Q-MyHC^+/-^ show reduced myofibrillar contractility^24, 25^, greater myofiber isometric tension development at submaximal Ca^2+^ levels^26^, and slower LV relaxation^27^.

In this study, we performed linked sn-ATAC-seq and sn-RNA-seq in R92W-TnT, R403Q-MyHC and littermate control mouse hearts at 5 weeks of age, with the goal of obtaining insights into transcriptional regulation of phenotypic heterogeneity in early-stage HCM. Our studies were performed in mouse heart tissue rather than human heart tissue, because HCM patients at early disease stage generally do not have clinical indications for cardiac biopsies or heart surgery. We hypothesized that a combination of mutation-specific changes in cardiomyocytes, and crosstalk between ventricular cardiomyocytes and fibroblasts, endothelial cells (ECs), vascular smooth muscle cells (VSMCs), macrophages underlie phenotypic variability in HCM, because mutant sarcomeric proteins are only expressed in cardiomyocytes. Our studies revealed activation of activator protein-1 (AP-1) transcription factors (TFs) and chromatin remodeler complexes (SWI/SNF, ISWI) in both mutant ventricular-cardiomyocytes. The transcriptional landscapes of mutant TnT and MyHC ventricular-cardiomyocytes were generally similar, whereas fibroblasts, macrophages, ECs, VSMCs were markedly different. Greater chromatin remodeling was seen in all cardiac cell types from mutant-TnT when compared to mutant-MyHC. Interrogation of cellular crosstalk using Cellchat predicted enrichment of pro-fibrotic signaling only in mutant-TnT, and pro-hypertrophic growth factor signaling to ventricular-cardiomyocytes in both mutants. Taken together, our studies provide an integrated view of transcriptional regulation in individual cardiac cell types, that contribute to phenotypic heterogeneity in HCM.

## RESULTS

### Study design

We performed deep cardiac phenotyping of mutant-TnT (R92W-TnT), MyHC (R403Q-MyHC) and littermate-control mice (CTR) at 5 weeks of age, which corresponds to early-disease stage. We examined cardiac morphology using echocardiography, histology, and isolated nuclei from the right and left ventricles (RV, LV), right and left atrial appendages (RAA, LAA) for ATAC-seq and RNA-seq. We compared gene expression and chromatin accessibility data between mutant and control mice using ArchR, SCENIC, Gene Ontology (GO), Ingenuity pathway analysis (IPA), KEGG, WikiPathways, CellChat (**Fig 1a-b**), to gain insights into transcriptional regulation and cellular crosstalk in cardiomyocytes and non-myocyte cells at early stage HCM.

**Figure 1.**
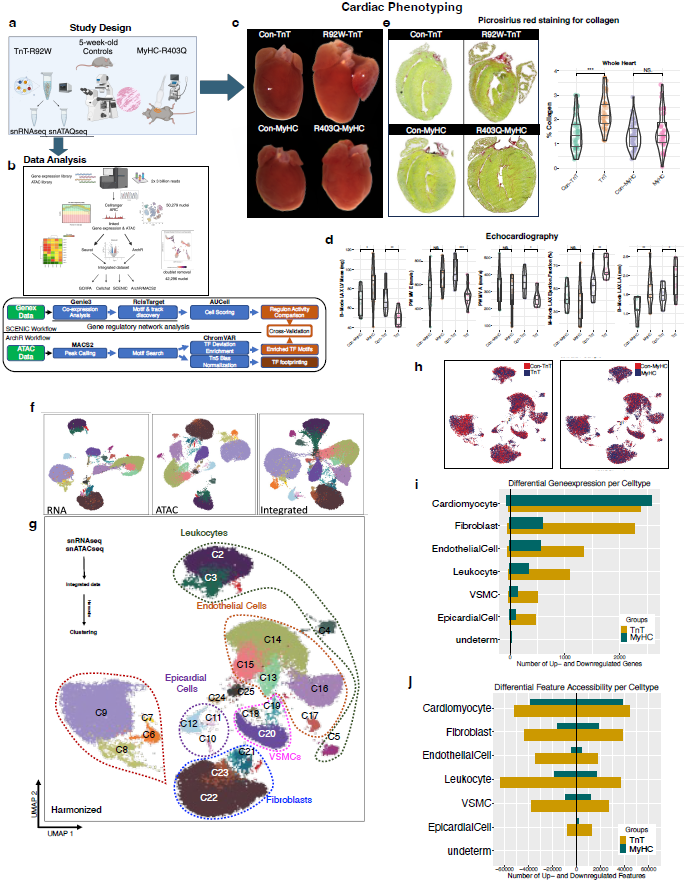
Study design, Cardiac phenotyping, Single nucleus data analysis. **a.** R92W-TnT and R403Q-MyHC mutant and littermate control mice were studied by echocardiography, histology, single-nuclei RNA-and ATAC-seq, at 5 weeks of age (early disease stage). Mutants were compared to littermate controls. **b.** Nuclei were isolated from the right and left ventricles (RV, LV) and atrial appendages (RAA, LAA). Single-nuclei gene-expression and ATAC libraries were obtained from the same nuclei, sequenced, and analyzed using Seurat, ArchR, SCENIC, CellChat. **c.** Cardiac morphology: Allele-specific cardiac remodeling was evident at early 5 weeks of age. Representative images show bi-atrial enlargement associated with smaller LV in TnT mutants and larger LV in MyHC mutants. **d.** Echocardiography: LV mass was significantly lower in TnT mutants and higher in MyHC mutants compared to littermate controls. Mitral E and A velocities, LV ejection fraction and LA diameter were significantly higher in TnT mutant. Welch’s t test: * p<05; ** p<0.01; *** p<0.001 **e.** Histology: Picrosirius red staining for collagen, revealed increased interstitial fibrosis only in TnT mutant hearts. Welch’s t test: * p<05; ** p<0.01; *** p<0.001 **f.** RNAseq and ATACseq data was integrated after dimensionality reduction, and harmonized. UMAP projection of all clusters based on RNAseq and ATACseq alone, following integration and harmonization, is shown. **g.** Cell identity of clusters obtained after integration and harmonization is presented. While most clusters could be assigned to a cell type, the identity of two clusters could not be clearly assigned. **h.** Experimental group identity is projected on the UMAP. All cell types could be identified in all groups. **i.** RNA-seq: Differential gene expression in TnT and MyHC mutants, compared to littermate controls for each cell type shows significant upregulation (adj. p<0.05) of the majority of differentially expressed genes (DEGs) in both mutants. TnT mutants have greater dysregulation of gene expression than MyHC mutants.**j.** ATAC-seq: Differences in chromatin accessibility are more pronounced in TnT mutant hearts, when compared to MyHC mutants. Non-myocyte cells show greater changes (FDR < 0.05) in chromatin accessibility than cardiomyocytes.

### Phenotypic heterogeneity at early disease stage

Mutant-TnT hearts demonstrated marked biatrial enlargement, whereas LA size was similar in MyHC-mutants, when compared to respective littermate controls (**Fig 1c**). Picrosirius staining revealed higher diffuse global fibrosis in TnT-mutants, but no difference in MyHC-mutants when compared to respective littermate-controls (**Fig 1d**). Echocardiography revealed lower left ventricular (LV) mass, mitral valve A velocity, and higher left ventricular ejection fraction (LVEF), left atrial (LA) size in mutant-TnT hearts compared to littermate-controls (**Fig 1e**). In contrast, MyHC-mutants had higher LV mass and LA diameter, similar LVEF, mitral A velocity size as littermate-controls (**Fig 1e**). Taken together histology and echocardiography reveal marked differences in cardiac phenotype at early disease stage, with interstitial fibrosis, LV diastolic dysfunction, LV hypercontractility without hypertrophy in TnT-mutants, and LVH with similar LV systolic/diastolic function in MyHC-mutants.

### Single nucleus gene expression and chromatin accessibility analysis in mutant hearts compared to littermate controls

To examine transcriptional regulation of phenotypic differences at early disease stage, we isolated nuclei from mutant and littermate-control hearts at 5 weeks of age, and performed linked sn-RNA-seq and ATAC-seq. We compared 23,304 nuclei from mutant hearts with 17,669 nuclei from controls, to identify cell-type and allele-specific transcriptional programs activated in early-stage HCM.

The RNA-seq and ATAC-seq data was integrated using Harmony, to generate 25 clusters that represent all known cardiac cell types (**Fig 1f-g**). Cluster identity was determined using known marker gene expression for each cell type, along with cluster-defining differential gene expression and chromatin accessibility (**Suppl. Fig 1a-c**). We identified clusters comprised of leukocytes (C1-5), cardiomyocytes (C6-9), epicardial cells (C10-12), endothelial cells (C13-17), vascular smooth muscle cells (C18-20) and fibroblasts (C21-23); two clusters could not be assigned to any cell type (**Fig 1g-h**).

As expected, ventricular-cardiomyocytes showed the highest differential gene expression between mutants and controls (**Fig 1i**). In all other cardiac cell types, mutant-TnT cells had greater dysregulation of gene expression, when compared to mutant-MyHC (**Fig 1i**). Additionally, greater chromatin remodeling was observed across all cardiac cell types in mutant-TnT, when compared to mutant-MyHC (**Fig 1j**). Taken together, our data indicates that at early disease stage, these two pathogenic variants in sarcomeric protein genes *Tnnt2* and *Myh6* are associated with profound changes in gene expression and chromatin accessibility in all cardiac cell types, with greater allele-specific differences in fibroblasts, leukocytes, endothelial cells (ECs), vascular smooth muscle cells (VSMCs) and epicardial cells when compared to cardiomyocytes.

### AP-1 transcription factors and chromatin remodeler complexes are activated in mutant ventricular cardiomyocytes at early disease stage

Unbiased clustering grouped cardiomyocytes into four clusters (C6-9), of which Cluster 9 (C9) was the largest (**Fig 2a**), and represented ventricular-cardiomyocytes, based on expression of genes such as *Tnnt2, Myh6* (**Suppl. Fig 1a-c**). Ventricular-cardiomyocytes demonstrated 1877 commonly upregulated genes between the 2 mutants, and a smaller proportion of upregulated genes specific to each mutant (n=228 in TnT; n=581 in MyHC); very few differentially expressed genes (DEGs) were downregulated (**Fig 2b**, **Suppl. Fig 2a-d**). A notable genotype-specific response was marked upregulation (log2FC >3) of the nucleus/golgi-localized zinc transporter *Slc39a11*, and the beta globin gene *Hbb-bs* in mutant TnT ventricular-cardiomyocytes (**Suppl. Fig 2b**). Gene Ontology (GO) analysis revealed enrichment for processes such as DNA transcription, post-translational protein modifications/breakdown in both mutant ventricular-cardiomyocytes (**Suppl. Fig 2g-h**). Additionally, IPA revealed that ventricular-cardiomyocytes from both mutants had signatures of cardiac hypertrophy signaling (**Fig 2c**), as well as upregulation of hypertrophy-associated genes (e.g. *Sorbs2, Pdlim5, Camk2d, Ryr2, Prkg1)* and TFs *(e.g. Mef2a/c, Gata4/6, Ep300)* (**Fig 2d**).

**Figure 2.**
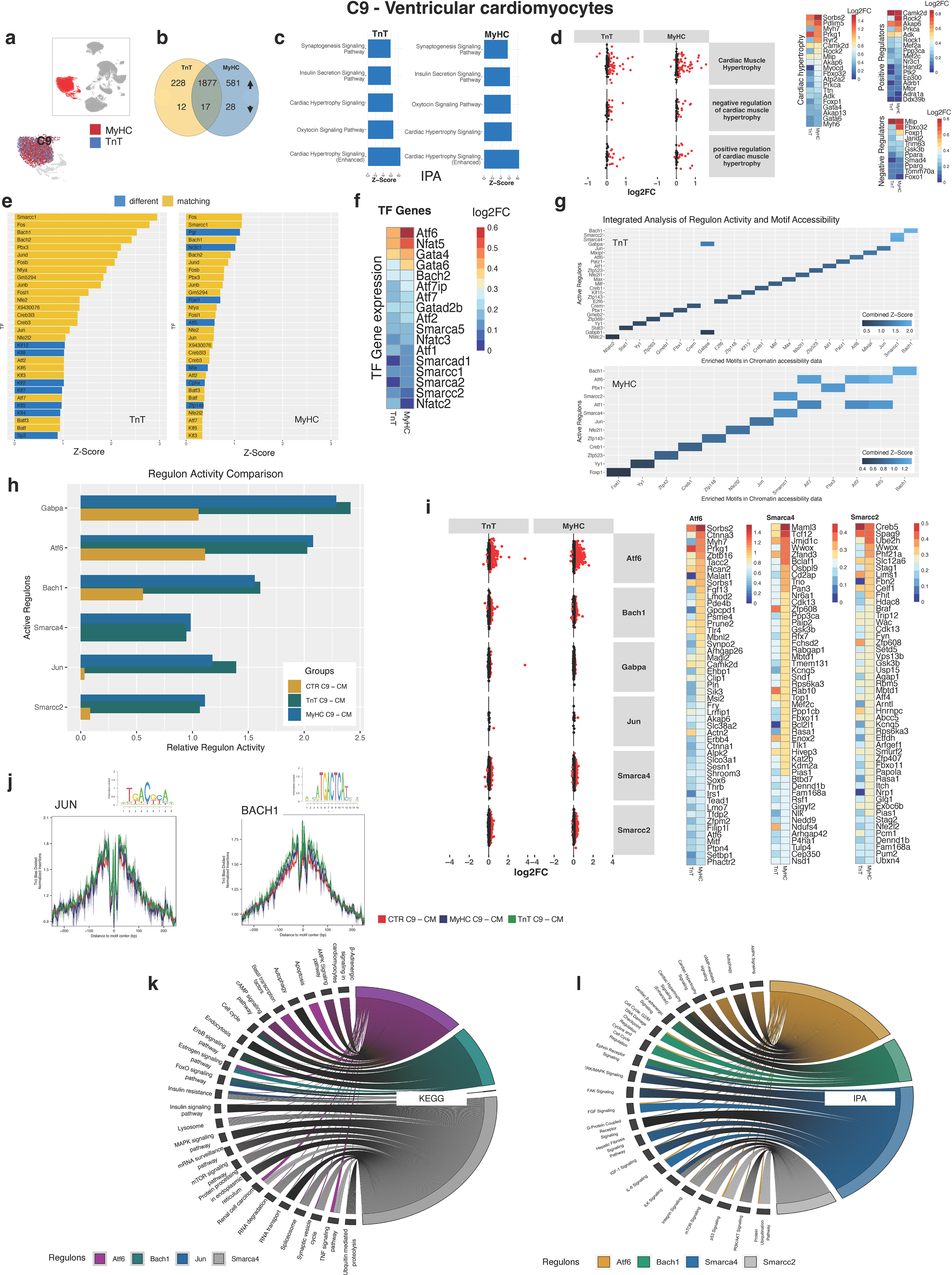
Ventricular cardiomyocytes (C9). **a.** ATAC-and RNA-seq: cell identity for each mutant in C9 (ventricular cardiomyocyte cluster) is presented. **b.** RNA-seq: Differentially-expressed genes (DEGs) in mutant cardiomyocytes were normalized to WT myocytes. Venn diagram shows majority of DEGs are upregulated in both mutant cardiomyocytes (adj. p<0.05). **c.** RNA-seq: Strong activation of cardiac hypertrophy signaling is predicted in both mutant cardiomyocytes by IPA, using all DEGs (adj. p<0.05); top 5 predicted pathways are presented. **d.** RNA-seq: Scatter plot and heat map of cardiac hypertrophy gene sets (AmiGO) in mutant MyHC cardiomyocytes compared to mutant TnT. (Red dots represent DEGs with adj. p<0.01, in cardiac hypertrophy gene sets). **e.** ATAC-seq: Bar graphs show the 30 most enriched TF motifs (FDR ≤ 0.05) in each mutant compared to WT, with yellow bars representing common motifs and blue bars distinct ones. Marked overlap (23/30) in TF motif enrichment in mutant cardiomyocytes, with higher z scores in mutant TnT. **f.** RNA-seq: Heatmap of differentially-expressed TF genes (adj p<0.05) identified by chromatin accessibility analysis, regulon analysis, GO pathway. **g.** Activated TF predicted by SCENIC and ArchR analysis: z-score for regulon activity and TF-motif-enrichment were aggregated (mean). TFs with high combined z-scores are presented. Bach1 and Smarcc1 have high combined z-scores (z ≥1.2) in both mutant ventricular cardiomyocytes.**h.** RNA-seq: Differential regulon activation in cardiomyocytes from mutants and WT. Smarcc and Jun are activated in both mutant cardiomyocytes, with very little activation in WT. **i.** RNA-seq: Scatter plot and heatmaps of regulon genes in mutant cardiomyocytes compared to WT. (Red dots represent differentially expressed regulon genes with adj p<0.01). Top 30 regulon genes are presented in the heat maps. **j.** ATAC-seq: Bulk TF footprinting across the mutant and WT genomes showed higher occupancy of Jun and Bach1 motifs in both mutant cardiomyocytes, compared to WT. **k.** RNA-seq: Top 10 processes enriched in differentially-expressed regulons genes, predicted by KEGG and IPA, by cross-referencing regulon genes with KEGG or IPA gene set lists. Matches with p<0.01 (Fisher exact test) were considered significant.

Chromatin accessibility was increased for 42,880 and 39,791 features, and decreased for 57,396 and 52,175 features, in mutant TnT and MyHC ventricular-cardiomyocytes, respectively (**Suppl. Fig 2e-f**). Motif enrichment analysis using ArchR predicted similar differential motif enrichment in the 2 mutants, with a stronger response in mutant-TnT when compared to mutant-MyHC (**Fig 2e**). Enrichment of AP-1 (activator protein 1) motifs^28^ was predicted in both mutants (**Fig 2e**), with stronger enrichment in mutant-TnT, whereas PGR and NR3C1 were only enriched in mutant-MyHC (**Fig 2e**). Analysis of TF gene expression confirmed modest upregulation of several subunits (*Smarcc1, Smarca2, Smarcc2, Smarca5, Smarcad1)* of the ATP-dependent chromatin remodeler complexes, SWI/SNF^29^, ISWI^30^, as well as *Gata4/6, Bach2,* and several members of the *Atf* and *Nfat* families of TFs (**Fig 2f**).

Next, in order to examine the association between increased chromatin accessibility and TF gene expression with downstream gene expression, we used SCENIC to identify groups of co-expressed genes (regulons) driven by a specific TF, and surveyed for matches between TF activation predicted by SCENIC (RNA-seq data) and ArchR (ATAC-seq data) by computing a combined z-score (mean of the respective z-scores), with the goal of strengthening our predictions of activated TFs (**Fig 2g**). The greatest differences between mutants and controls were seen for Smarcc2, Smarca4, Jun, Bach1 regulons (using RNA-seq data), with controls showing no Smarca4 activity and very small Jun and Smarcc2 activity (**Fig 2 g-i**). Pseudo bulked TF-footprinting of the JUN and FOS motifs showed higher Tn5 biased normalized insertions reflecting higher TF occupancy in mutants compared to controls, with mutant-TnT ventricular-cardiomyocytes demonstrating greater occupation than mutant-MyHC (**Fig 2j**). Smarca4, Jun, Bach1, Atf6 regulon genes were enriched for biologic processes such as autophagy, protein processing in ER, ubiquitin mediated proteolysis, lysosome, apoptosis (**Fig 2k**), and signaling pathways such as ERK/MAPK, IGF1, integrin, FAK, mTOR, beta adrenergic and cardiac hypertrophy signaling. (**Fig 2l**).

In summary, our analysis of linked RNA-seq and ATAC-seq at early disease stage suggest activation of cardiac hypertrophy signaling, AP-1 TFs and chromatin remodeler complexes in both mutant ventricular-cardiomyocytes. We hypothesize that these pathogenic variants in *Tnnt2* and *Myh6* lead to activation of signaling pathways and AP-1 TFs (**Suppl. Fig 3a**), which recruit chromatin remodeler complexes, SWI/SNF, ISWI (**Suppl. Fig 3b**). Chromatin remodeling enables activation of late response genes involved in autophagy, proteostasis, redox, mitochondrial metabolism, and facilitates CAMKII-induced DNA binding of Nfat/Mef2/Gata4 to promote myocyte hypertrophy^31^ at early disease stage (**Suppl. Fig 3c**). Activation of the glucocorticoid receptor which is only seen in mutant-MyHC cardiomyocytes could potentiate myocyte hypertrophy in MyHC-mutant hearts (**Suppl. Fig 3d**).

### Profibrotic gene programs are selectively activated in mutant TnT fibroblasts at early disease stage

Iterative clustering identified three cardiac fibroblast subpopulations (C21-23). Cluster C22 (**Fig 3a**) was the largest (7811 nuclei) and showed the highest expression of activated fibroblast marker genes (e.g. C*ol8a1, Postn, Col3a1, Col1a1, Thbs1*), but only in mutant-TnT fibroblasts (**Suppl. Fig 4a, b**). Hence, we focused our analysis on this fibroblast cluster. Fibroblasts in the C22 cluster showed 531 commonly upregulated genes in both mutants; 1455 genes were only upregulated in mutant-TnT fibroblasts, and 25 genes were only upregulated in mutant-MyHC; very few genes were downregulated (**Fig 3b**, **Suppl Fig 4c-f**). Mutant-TnT fibroblasts had signatures of activated fibrosis signaling by IPA (**Fig 3c**), and upregulation of several genes (e.g. *Col3a1, Col1a2, Tgfbr1-3, Smad 3-4, Tcf4*) and TFs (e.g. *Ep300, Foxp2, Crebbp, Tcf7l1/2*), associated with pro-fibrotic TGFβ and Wnt signaling pathways (**Fig 3d, e**).

**Figure 3.**
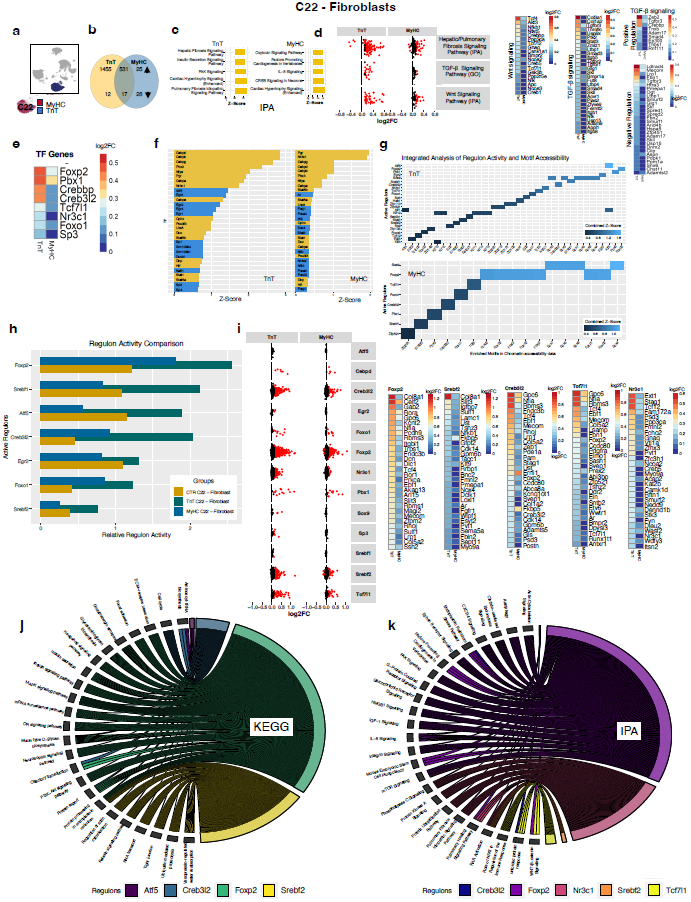
Cardiac fibroblasts (C22). **a.** Three clusters were identified as fibroblasts, of which C22 was the largest. **b.** RNA-seq: Greater dysregulation of gene expression in mutant TnT fibroblasts, when compared to mutant MyHC. Most dysregulated genes are upregulated in fibroblasts from both mutants (adj p< 0.05). **c.** RNA-seq: Top 5 signaling pathways predicted to be activated by ingenuity pathway analysis (IPA) using all differentially expressed genes (DEGs) with adj. p<0.05. Activation of fibrosis signaling was only predicted in mutant TnT fibroblasts. **d.** RNA-seq: Analysis of IPA and GO gene lists for profibrotic signaling pathways Wnt and TGF-β, revealed greater dysregulation of gene expression in mutant TnT fibroblasts, when compared to MyHC mutants. (Red dots represent DEGs with adj. p<0.01). All DEGs in Wnt and TGFβ signaling pathways are presented in the heat maps. **e.** RNA-seq: Heatmap of differentially-expressed TF genes (adj. p<0.05) identified by chromatin accessibility analysis, regulon analysis, and previously described to be involved in fibrosis. Foxp2, Pbx1, Crebbp and Creb3l2 are the top 4 differentially-expressed TF genes in fibroblasts from both mutants. **f.** ATAC-seq: Bar graphs shows the 30 most enriched TF motifs (FDR ≤ 0.05) in mutant fibroblasts compared to controls. Mutant TnT fibroblasts show greater chromatin remodeling, when compared to mutant MyHC. Agreement (yellow bars) between the 2 mutants was observed for 14/30 motifs. **g.** RNA-seq and ATAC-seq: Integration of motif accessibility and the regulon activity showing a combined (average) z-score. ATF5 and FOXP2 had the highest z-score (z >1.2) in mutant TnT fibroblasts, and several members of the SOX and FOX family of TFs had the highest z-score (z >1) in mutant MyHC fibroblasts. **h.** RNA-seq: Relative regulon activity in mutant and littermate control fibroblasts show stronger TF activation in mutant TnT fibroblasts. **i.** RNA-seq: Scatter plot and heatmaps of regulon genes in mutant fibroblasts compared to controls. (Red dots represent differentially-expressed regulon genes with adj p<0.01). Top 30 regulon genes are presented in the heat maps. **j, k.** RNA-seq: Biologic processes associated with differentially-expressed regulon genes in fibroblasts from TnT and MyHC mutants, predicted by KEGG and IPA; band thickness reflects number of matching genes.

Analysis of C22 ATAC-seq data revealed increased chromatin accessibility for 40,567 and 21,287 features, and decreased accessibility for 58,030 and 21,287 features in mutant TnT and MyHC fibroblasts, respectively (**Suppl Fig 4g-h**). Motif enrichment analysis using ArchR predicted enrichment for several CEBP family members, PGR and NR3C1 in both mutants, with a stronger response in mutant-TnT (**Fig 3f**). The greatest differences between both mutants and controls (by SCENIC and ArchR analyses) were observed for Foxp2, Tcf7l1, Creb3l2, Srebf2 regulons (**Fig 3g-i**). These regulons are enriched for biologic processes such as focal adhesion, glycosaminoglycan synthesis, protein synthesis/ER processing/degradation (**Fig 3j**), and signaling pathways such as MAPK, PI3K-AKT, IGF1, Wnt-β catenin signaling, actin cytoskeletal signaling, FAK signaling, ER stress pathway (**Fig 3k**).

This data led us to hypothesize that activation of canonical TGFβ signaling via SMADs, and Wnt-β catenin signaling (**Fig 3c-d**) by P38-MAPK activation (**Fig 3j-k**) promotes mutant-TnT fibroblast activation. Upregulation of collagen gene expression leads to increase in collagen synthesis which activates ER-localized TFs^32^, SREBF2 and CREB3L2 (**Fig 3h-i**), leading to increased collagen secretion and interstitial fibrosis in mutant-TnT hearts (**Fig 1e**).

### Mutant TnT endothelial cells have greater transcriptional dysregulation than mutant MyHC

Unbiased clustering identified five endothelial cell (EC) clusters (C13-C17) with C14 being the largest EC cluster with 6770 nuclei (**Fig 4a**) and likely representing arterial ECs – hence we focused our analysis on C14. We observed 292 commonly upregulated genes in both mutants; 297 genes were only upregulated in mutant-TnT, and 51 genes were only upregulated in mutant-MyHC ECs; very few genes were downregulated (**Fig 4b**, **Suppl Fig 5a-d)**. Both mutant-ECs had signatures of integrin-mediated cell adhesion and focal adhesion, whereas only mutant-TnT ECs showed strong activation of EGFR1 and MAPK signaling pathways (**Fig 4c**) and associated genes (**Fig 4d**).

**Figure 4.**
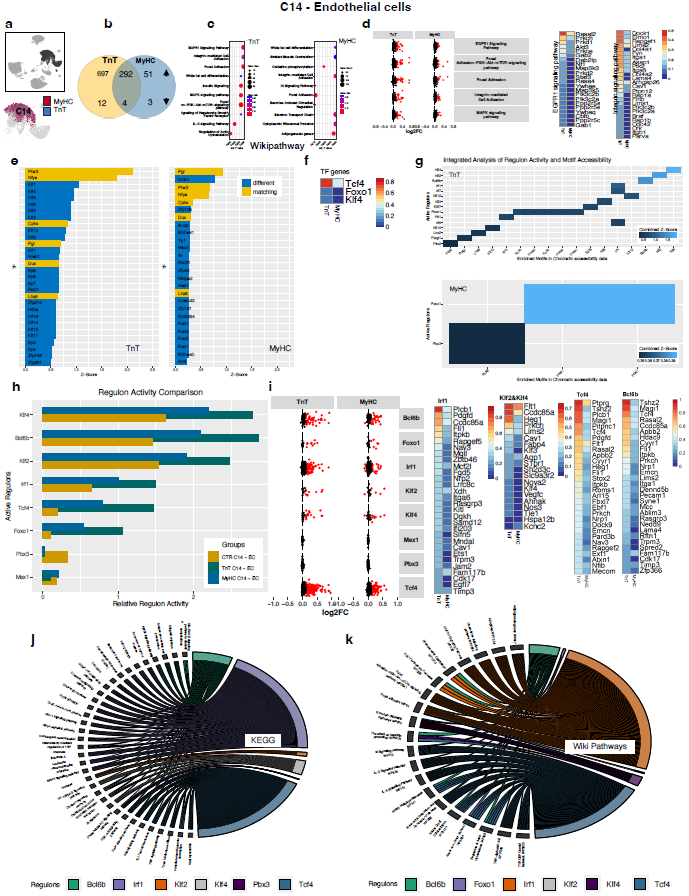
Endothelial cells (ECs, C14). **a.** RNA-, ATAC-seq: Five EC clusters with C14 being the largest. **b.** RNA-seq: Greater dysregulation (adj. p<0.05) of genes in mutant TnT, when compared to mutant MyHC; most dysregulated genes are upregulated. **c.** RNA-seq: Top 10 activated signaling pathways in ECs, predicted by Wikipathway GSEA, using upregulated genes (adj. p<0.05). EGFR1 and MAPK signaling activation is only predicted in mutant TnT ECs, whereas focal adhesion and integrin-mediated cell adhesion is predicted in both mutants. **d.** RNA-seq: Scatter-plot and heatmaps for genes in signaling pathways, predicted by Wikipathways. Mutant TnT ECs had greater dysregulation of genes involved in EGFR1, MAPK, focal adhesion and integrin-mediated signaling, when compared to mutant MyHC. (Red dots represent genes with adj. p<0.01) All DEGs in EGFR1 and integrin signaling pathways are presented in the heat maps. **e.** ATAC-seq: Bar graphs shows the 30 most enriched TF motifs (FDR ≤ 0.05) in mutant ECs compared to WT. Differential TF motif enrichment shows marked differences between mutant ECs, with only 6/30 matches (PBX3, NFYA, PGR, CPHX, DUX, LHX3); several members of the KLF family of TFs were only enriched in mutant TnT ECs. **f.** RNA-seq: Differential expression (adj. p<0.05) of TF genes identified by motif enrichment analysis shows higher expression of *Tcf4* in both mutant ECs; *Foxo1, Klf4* expression was only increased in mutant TnT ECs. **g.** RNA-seq and ATAC-seq: Integration of motif accessibility and regulon activity by combined (average) z-score shows z >1.5 for Klf2, Klf4 in mutant TnT ECs; no regulons with z >1.0 were identified in mutant MyHC. **h.** RNA-seq: Differential regulon activity analysis shows highest activity for Klf2, Klf4, Bcl6b, Irf1, Tcf4, Foxo1 regulons in mutant TnT ECs. **i.** RNA-seq: Scatter-plot and heat maps of regulon genes shows greater upregulation of genes for Bcl6b, Irf1, Tcf4 in TnT mutant ECs, when compared to MyHC mutants. (Red dots represent differentially expressed regulon genes with adj p<0.01). Heat maps show top 30 differentially-expressed regulon genes. **j,k.** RNA-seq: Biologic processes associated with regulon genes were identified by cross-referencing with KEGG and Wikipathways gene set lists. Matches with p<0.01 were considered significant.

Analysis of C14 ATAC-seq data revealed increased accessibility for 22,526 and 9,436 features, and decreased accessibility for 57,277 and 13,599 features for mutant TnT and MyHC ECs respectively (**Suppl. Fig 5e-f**). Analysis of differential motif enrichment using ArchR revealed marked differences between the mutant ECs, with an overall stronger response in TnT-mutants, and enrichment for KLF, SP and ZFP family of TFs only in mutant-TnT ECs (**Fig 4e**). Analysis of differentially-expressed TF genes (**Fig 4f**) revealed modest upregulation of *Klf4, Foxo1* only in mutant-TnT ECs, and *Tcf4* in both mutants. The greatest differences between the study groups, identified by SCENIC and ArchR analysis, were observed for Tcf4, Foxo1, Irf1, Klf2, Klf4, Bcl6b regulons (**Fig 4g-i**). These regulons are enriched for biologic processes such as cell adhesion, cell cycle (**Fig 4j**). and signaling pathways such as EGFR1, focal adhesion-PI3K-Akt-mTOR, Ras and MAPK (**Fig 4k**).

In summary, mutant-TnT ECs demonstrate greater dysregulation of gene expression and chromatin accessibility than mutant-MyHC ECs. Our analysis of RNA-seq and ATAC-seq data in ECs leads us to hypothesize that hyperdynamic LV function in TnT-mutants increases shear^33^ and circumferential^34^ stress in coronaries (**Suppl. Fig 6a**), which activates integrins^35^ and kinases such as focal adhesion kinase (FAK) – this leads to EGFR transactivation^36^ (**Fig 4c-d**), which induces Irf1 (via STAT)^37^ and Klf2/4 (via ERK, MEF2 phosphorylation)^38^ (**Fig 4e-i**, **Suppl. Fig 6b**). Klf2/4 stimulate NOS-3 (nitric oxide synthase-3) and nitric oxide (NO) generation and inhibit EC activation/proliferation. Furthermore, activation of Irf1 can inhibit angiogenesis^39^. Taken together, this data could explain our clinical perfusion PET-imaging results of high myocardial blood flow at rest in HCM patients^3^, and pathology studies suggesting mismatch between myocardial mass and coronary vasculature^2^ (which contributes to microvascular ischemia and angina^3^ in HCM).

### Mutant TnT vascular smooth muscle cells have greater dysregulation of gene expression and chromatin remodeling when compared to mutant MyHC

Unbiased clustering identified three vascular smooth muscle (VSMC) clusters, C18-20. We analyzed the largest cluster, C20 (**Fig 5a**) with 2622 nuclei. Similar to other cell types, we observed greater dysregulation of gene expression in mutant-TnT, when compared to mutant-MyHC VSMCs (**Fig 5b-c**). The DEGs in mutant-TnT VSMCs were enriched for biologic processes such as focal adhesion, inositol mediated signaling, regulation of EC migration, whereas mutant-MyHC ECs showed enrichment for morphogenesis and filament sliding (**Fig 5d**). Ingenuity pathway analysis (IPA) predicted activation of fibrosis signaling in both mutants, and NO signaling only in mutant-TnT VSMCs (**Fig 5e**). Modest upregulation of a few genes in the PDGF signaling pathway (GO) was seen, with greater changes in mutant-TnT when compared to mutant-MyHC VSMCs (**Fig 5f**).

**Figure 5.**
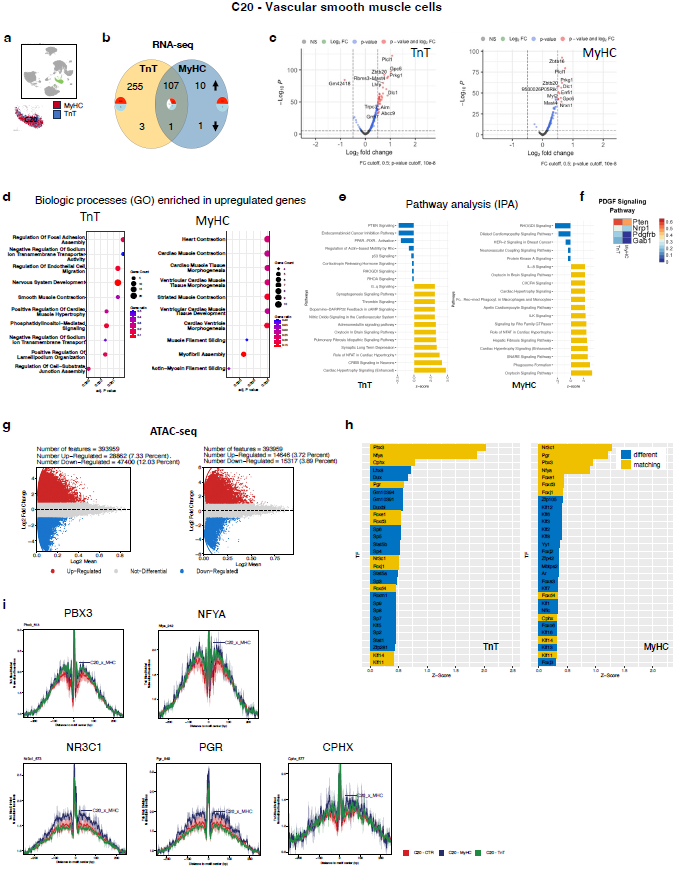
Vascular smooth muscle cells. **a.** Three clusters were identified as VSMCs, with C20 being the largest. **b.** RNA-seq: Differentially expressed genes (adj p<0.05) in mutants normalized to WT shows greater numbers of upregulated genes in mutant TnT VSMCs when compared to mutant MyHC. **c.** RNA-seq: Volcano plot of DEGs in both mutants compared to WT. **d.** RNA-seq: Top 10 GO biologic processes enriched in mutant VSMCs using all DEGs include ‘regulation of focal adhesion’, ‘endothelial cell migration’ and ‘smooth muscle contraction’ only in mutant TnT VSMCs. **e.** RNA-seq: IPA analysis of all DEGs predicts activation of NO signaling in mutant TnT ECs, and fibrosis signaling in both mutants. **f.** RNA-seq: Four genes in the PDGF signaling pathway are dysregulated in mutants. **g.** ATAC-seq: Mutant TnT VSMCs have greater numbers of differentially-accessible features when compared to mutant MyHC. **h.** ATAC-seq: Bar graph shows the 30 most enriched (FDR ≤ 0.05) TF motifs. Yellow bars represent common motifs and blue bars distinct ones: 11/30 TF motifs (including PBX3, NYFA, NR3C1, PGR) are enriched in both mutant VSMCs. **i.** ATAC-seq: Bulk TF footprinting across the mutant and control genomes showed increased occupancy of PBX3, NYFA in both mutant VSMCs, and NR3C1, PGR, CPHX motifs only in mutant MyHC.

Analysis of ATACs-seq data revealed that 28,862 and 14,646 features were more accessible, whereas 47,400 and 15,317 were less accessible in the mutant TnT and MyHC VSMCs respectively (**Fig 5g**). TF motif enrichment analysis predicted marked differences between the mutants with agreement only for PBX3, NFYA, CPHX, and PGR (**Fig 5h**). Since the C20-VSMC cluster is small, regulon analysis could not be carried out reliably. TF-footprinting revealed increased binding for CPHX, PBX3, NYFA motifs in both mutant VSMCs, and stronger binding for NR3C1, PGR motifs only in mutant-MyHC VSMCs (**Fig 5i**).

### Cardiac macrophages in mutant TnT demonstrate greater transcriptional dysregulation than mutant MyHC

Unbiased clustering identified 5 leukocyte clusters (C1-5), of which C2 was the largest (3657 nuclei) and represented macrophages (**Fig 6a**). We found 294 commonly upregulated genes in both mutants; 580 genes were only upregulated in mutant-TnT macrophages, and 18 genes were only upregulated in mutant-MyHC; very few genes were downregulated (**Fig 6b**, **Suppl Fig 7a-d**). Gene ontology analysis revealed that DEGs in both mutants were enriched for biologic processes related to the actin cytoskeleton, endocytic vesicle, intracellular membrane bound organelle (**Fig 6c**); upregulation of genes involved in phagosome formation/maturation and phagocytosis was more prominent in mutant-TnT macrophages, when compared to mutant-MyHC (**Fig 6d**).

**Figure 6.**
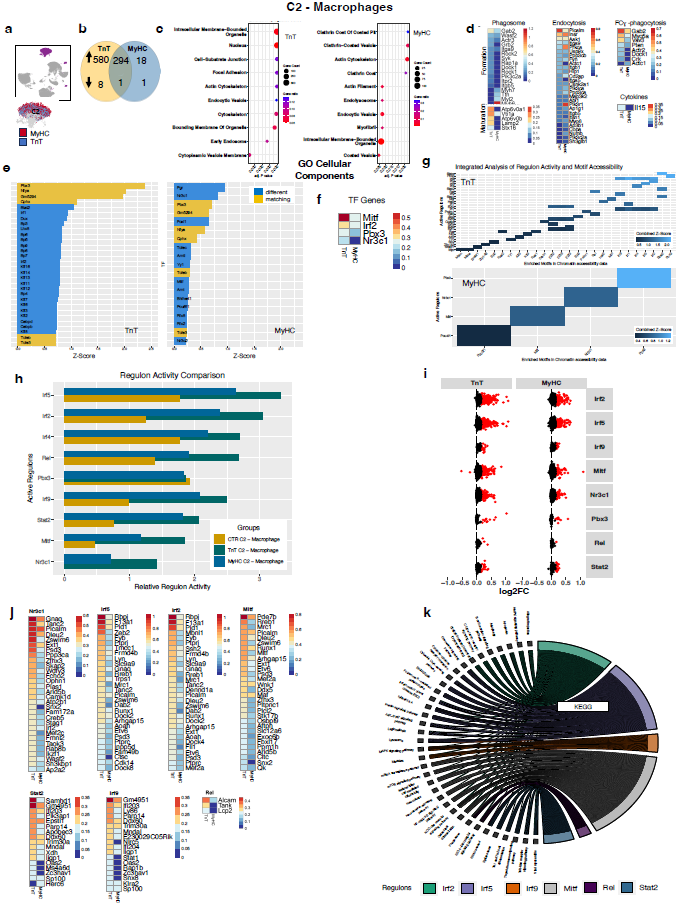
Cardiac macrophages (C2). **a.** RNA-and ATAC-seq: C2 (cardiac macrophages) is the largest leukocyte cluster. **b.** RNA-seq: Mutant TnT macrophages show greater dysregulation (adj p<0.05) of gene expression than mutant MyHC, with most dysregulated genes being upregulated. **c.** RNA-seq: Top 10 pathways predicted to be enriched by GO Cellular Component pathway analysis using all DEGs. Activation of endocytosis and phagocytosis was predicted in both mutants, with more genes matching in the TnT mutant. **d.** RNA-seq: Heatmap for DEGs for phagosome formation, maturation, endocytosis, FCLJ receptor-mediated phagocytosis show greater upregulation in mutant TnT. Cytokine gene expression analysis showed a small statistically significant upregulation of IL-15 gene expression in mutant TnT macrophages. (Red dots represent DEGs with adj p<0.01); all DEGs in GO genesets for phagosome formation, maturation, endocytosis, FclZl phagocytosis, cytokine genes are presented in the heat maps. **e.** ATAC-seq: Top 30 differentially-accessible TF motifs (FDR ≤ 0.05) in mutants compared to WT. Marked differences are seen between mutants; 6/30 TF motifs, including PBX3, NFYA, CPHX showed higher accessibility in both mutants, with greater changes in mutant TnT when compared to mutant MyHC. **f.** RNA-seq: Heatmap of differentially-expressed TF genes (adj p<0.05) identified by chromatin accessibility analysis, regulon analysis shows greater activation in mutant TnT when compared to mutant MyHC. **g.** RNA-seq and ATAC-seq: Integration of motif accessibility and regulon activity showing the combined (average) z-score shows z >1.2 for Pbx3 in both mutants; Stat2 and several Irf family members were only seen in mutant TnT. **h.** RNA-seq: Regulon activity analysis shows activation of several Irf family members (Irf2, 4, 5, 9), Stat2, Rel, Nr3c1, Mitf in both mutants with higher z-score in mutant TnT. **i, j.** RNA-seq: Scatter-plot and heatmaps of regulon genes in both mutants show greater upregulation in mutant TnT. (Red dots represent differentially-expressed regulon genes with adj p<0.01). Top 30 differentially expressed regulon genes are presented in the heat maps. **k.** RNA-seq: Regulon-Geneset comparison using KEGG with strength of interaction shown by number of matched genes shows that Irf5 is most influential for endocytosis and FC gamma receptor-mediated phagocytosis.

Analysis of ATAC-seq data revealed increased accessibility of 39,146 and 24,852 features, and decreased accessibility of 92,653 and 37,637 features for TnT and MyHC mutants respectively (**Suppl Fig. 7e-f**). Analysis of differential motif enrichment using ArchR revealed marked differences between the 2 mutants, with modest enrichment for KLF, SP, STAT2, CEBP family of TFs only in TnT-mutant macrophages (**Fig 6e**). Analysis of TF gene expression revealed modest upregulation of *Mitf, Irf2, Pbx3* in both mutants, with mutant-TnT macrophages demonstrating greater upregulation than mutant-MyHC. Regulons with the greatest differences between mutants and controls were observed for Irf 2/4/5/9, Nr3c1, Stat2 and Mitf (**Fig 6g-j**), which were enriched for biologic processes such as endocytosis, phagocytosis, cytosolic DNA sensing pathway, phagosome, and signaling pathways such as RIG/NOD-like receptor, NF*k*b, JAK-STAT, chemokine signaling pathways (**Fig 6k**).

In summary, analysis of RNA-seq and ATAC-seq data led us to hypothesize that resident macrophages in mutant hearts have a transcriptional profile that is closer to M1 polarization than M2, with mutant-TnT macrophages showing greater changes than mutant-MyHC (**Suppl. Fig 8a**). Based on our previous data showing mitochondrial dysfunction in TnT-mutants at early disease stage^15^, this data suggests that macrophages could play an important role in clearance of damaged mitochondria and other debris from ventricular cardiomyocytes^40^ (**Suppl. Fig 8b**). Damaged organelles (DAMPs) are recognized by macrophage pattern recognition receptors, leading to upregulation of NF-κB and IRF5 which are pro-inflammatory^41^. In this study, we did not observe significant upregulation of cytokine gene expression (**Fig 6D**), which we attribute to activation of the glucocorticoid receptor NR3C1, which can block the pro-inflammatory effects^42^ of NFκB and IRF activation (**Suppl. Fig 8c**).

### Cell-Cell communication Analysis

Mutant sarcomeric proteins are only expressed in cardiac myocytes, but all cardiac cell types demonstrate chromatin remodeling and changes in gene expression at early disease stage, suggesting the role of paracrine signaling. We used CellChat to examine intercellular communications and predict autocrine/paracrine signaling in mutant and control hearts. Cellchat predicts intercellular communication and signaling pathways, based on the expression of ligand-receptor pairs. Our analysis of RNA-seq data using CellChat revealed an overall increase in intercellular communications in both mutant hearts when compared to controls, with mutant-TnT demonstrating the greatest changes (**Fig 7a**). These changes were due to stronger interactions between ventricular cardiomyocytes (C9), ECs (C14, C16) and fibroblasts (C21, C22) in mutant hearts, when compared to controls (**Suppl Fig 10a)**. The mutant-TnT heart was highly enriched for ephrin B (which can influence diastolic function^43^), and pro-fibrotic periostin, TGFβ signaling, whereas mutant-MyHC showed marked enrichment for pro-hypertrophic^44^ neuregulin signaling (**Fig 7b-c**). PDGF and ANGPT signaling were predicted to be stronger in mutant-TnT, whereas VEGF signaling was similar in both mutants (**Fig 7b**).

**Figure 7.**
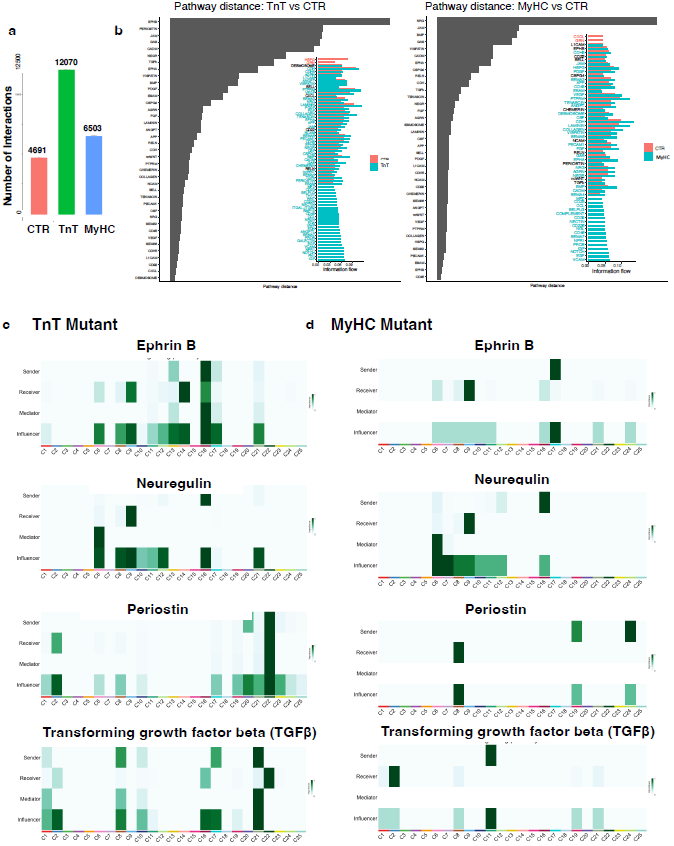
Prediction of cellular crosstalk by CellChat. Cell Chat uses RNA-seq data to infer ligand-receptor-pairings between different clusters which reflects paracrine and autocrine signaling. **a.** Higher numbers of interactions are predicted in cells from mutant TnT (12070) and mutant MyHC (6503), when compared to controls (4691). **b.** Pathway distance plot (grey bars) shows the difference in pathway activity between mutants and controls. Bar length reflects the degree of difference in the pathway activity between the two, with longer bars reflecting greater difference in pathway activity. The adjacent plot shows direct comparison of the information flow in each mutant, compared to controls. **c.** Cell-Cell-Signaling for selected pathways. Darker green indicates greater importance. Sender expresses the ligand and receivers express the receptor. Mediators and receivers influence the signaling but are not directly involved in signaling.

Next, we assessed disease-specific intercellular communications that are only identified in one or both mutants and not present in controls (**Suppl Fig 9b-e**). Upregulation of several ligand-receptor pairs/signaling, including NPR1, VCAM, NOTCH, EGF, IGF were only present in mutant hearts. Furthermore, only mutant-TnT showed expression of galectin, and pro-fibrotic ligands/signaling, including FN1, EDA, Wnt, ITGB2, BMP10, EDN (**Suppl Fig 9b-e**).

Analysis of outgoing communication from cardiomyocytes was performed to assess genotype-specific cellular crosstalk: antihypertrophic^45^ natriuretic peptide receptor 1 (NPR1) and vascular adhesion molecule (VCAM) were specific to mutant myocytes and not seen in controls (**Suppl Fig 10a-b**), whereas cyclic GMP-AMP synthase (GAS) was specific to mutant-TnT myocytes (**Suppl Fig 10c**, **Fig 7b**), and Agrin (AGRN) was specific to mutant-MyHC myocytes (**Suppl Fig 10d**). Interestingly, endothelin (EDN) signaling was only predicted to be activated in mutant-TnT, with the fibroblast cluster C21 as the source, and cardiomyocytes (C6-9) as recipients (**Suppl. Fig 10e**).

Taken together, our analysis of cell-cell communication in mutant TnT and MyHC hearts at early disease stage, reveals similar dysregulation of signaling pathways in mutant myocytes, and selective activation of pro-fibrotic pathways in mutant-TnT.

## DISCUSSION

We examined transcriptional regulation of cardiac phenotypic heterogeneity in early stage HCM by analysis of single nuclei isolated from 2 HCM mouse models with distinct cardiac phenotypes, using linked snRNA-seq and snATAC-seq. Since mutant sarcomeric proteins are only expressed in cardiomyocytes, we hypothesized that these pathogenic variants in *Tnnt2* and *Myh6* induce large, genotype-specific transcriptional changes in cardiomyocytes, as well as paracrine signaling from ventricular cardiomyocytes, that lead to differences in LV mass, cardiac mechanics and fibrosis at early disease stage. Our multiome analysis revealed similar transcriptional landscapes and activation of chromatin remodeler complexes (SWI/SNF, ISWI) in mutant ventricular cardiomyocytes and marked differences in gene expression and chromatin remodeling in cardiac fibroblasts, macrophages ECs, VSMCs, at early disease stage, suggesting an important role for non-myocyte cells in phenotypic heterogeneity in HCM. Contrary to our original hypothesis, ventricular cardiomyocytes were not predicted to drive dysregulated signaling pathways in fibroblasts, ECs or VSMCs.

### Cardiac HCM phenotype

The pathologic hallmarks of HCM are myocyte hypertrophy, fibrosis, microvascular remodeling. Clinically, HCM patients are classified into 3 hemodynamic groups, namely, obstructive, non-obstructive and labile obstructive, based on the presence/absence of LV obstruction at rest/provocation^46^. In each hemodynamic group, the location and extent of cardiac hypertrophy, fibrosis, coronary microvascular dysfunction is very variable, even in individuals expressing the same pathogenic variant. But very little is known about the molecular mechanisms driving phenotypic variability in human HCM, because of lack of availability of human heart tissue at early disease stage.

### Myocyte Ca^2+^ handling can influence cardiac HCM phenotype

Abnormalities in myocyte Ca^2+^ handling are common in HCM, and can lead to changes in systolic/diastolic function, as well as activation of pro-hypertrophic calcium-calmodulin kinase II (CAMKII) signaling (**Suppl Fig 3c**). Our analysis of snRNA-seq data revealed upregulation of several genes implicated in Ca^2+^ signaling^47^ (*Camk2d, Ryr2, Prkca*) in both mutant ventricular cardiomyocytes, which could promote myocyte hypertrophy^47^ (**Fig 2d**). Prior studies in R403Q-MyHC mutant-myocytes demonstrate reduced SR-Ca^2+^ stores without changes in diastolic Ca^2+^ or Ca^2+^ transients at 4 weeks of age^48^, which could explain our echocardiography results of similar LV systolic and diastolic function at 5 weeks of age. In contrast, the R92W-TnT variant causes increased myofilament Ca^2+^sensitivity, which is predicted to increase myofilament Ca^2+^binding, resulting in greater cytosolic Ca^2+^buffering, lower peak systolic Ca^2+^, higher peak tension, slower myofilament Ca^2+^dissociation, higher diastolic calcium^49^, positive inotropy and negative lusitropy^49^. These computer modeling results are confirmed by prior Ca^2+^ transient measurements in isolated R92W-TnT myocytes^50^ and could explain our echocardiography results of higher LVEF and diastolic dysfunction, in mutant-TnT hearts. Interestingly, cardiac myocytes isolated from R92W-TnT hearts have been reported to exhibit shorter baseline sarcomere lengths than controls, and patients expressing this pathogenic variant have mild or no cardiac hypertrophy at early disease stage. At the cellular level, increase in cardiomyocyte size is thought to be the main contributor to increase in LV mass^51^, because adult cardiomyocytes comprise ∼70–80% of heart mass and have little mitotic potential. Hence, smaller baseline mutant myocyte size in R92W-TnT, which has been attributed to higher basal Ca^2+^-associated activation, likely contributes to smaller heart size observed in TnT-mutant mice despite activation of hypertrophic signaling pathways and TFs.

### Myocyte hypertrophy

Since myocyte hypertrophy is a hallmark of HCM, we used our integrated snATAC-seq and snRNA-seq data from ventricular cardiomyocytes, to gain insights into the transcriptional regulation of LVH in early stage HCM. We identified large transcriptional changes in ventricular cardiomyocytes, and pathway analysis predicted activation of cardiac hypertrophy in both mutants (**Fig 2c**). Both mutant ventricular-cardiomyocytes demonstrated upregulation of genes encoding several pro-hypertrophic TFs (**Fig 2f**) and kinases such as *Camk2d, Rock2, Prkca, Akap6*. (**Fig 2 d, f**). Cellchat analysis of RNA-seq data predicted incoming pro-hypertrophic IGF signaling (from C1/C2) only in mutant ventricular-cardiomyocytes, which could activate MAPK signaling and induce transcription of early response genes *Fos, Jun* (**Suppl Fig 3b**). Induced Fos and Jun proteins (AP-1 TFs)^52^ have been demonstrated to function as pioneer TFs by binding nucleosomal DNA, and recruiting the chromatin remodeler complexes (SWI/SNF, ISWI)^29, 51^ to open chromatin and trigger transcription of late response genes that lead to the molecular cardiac phenotype (characterized by activation of the fetal gene program^51^, changes in mitochondrial function, metabolism, redox). Mammalian SWI/SNF complexes regulate genomic architecture by using energy from ATP hydrolysis to disrupt contact between DNA and histones, leading to nucleosome disassembly^29^. The mammalian SWI/SNF complex consists of 4 core subunits: one of 2 ATPase subunits, BRM (*Smarca2*) or BRG1 (*Smarca4*), BAF155 (*Smarcc1*), BAF170 (*Smarcc2*) and INI1/SNF5/BAF47 (*Smarcb1*)^53^. Our analysis of RNA-seq, ATAC-seq data, TF-footprinting, regulon analysis (**Fig 2e-j**) support activation of AP-1 TFs and chromatin remodeler complexes (**Fig 2e-h**) in early stage HCM. Prior studies in experimental models^51, 54^ reveal the importance of BRG1 (*Smarca4*) in pathologic cardiac hypertrophy^54^. BRG1 (*Smarca4*) is silenced in adult cardiac myocytes, but reactivated by cardiac stresses such as increased afterload resulting from TAC (transverse aortic constriction) or hypertension^54^, leading to the formation of SWI/SNF complexes that remodel chromatin, activate the fetal cardiac gene program^54^ and promote cardiac remodeling. An immunohistochemical study of BRG1 expression in human heart tissue revealed higher BRG1 expression in HCM than DCM, and other causes of LVH^55^. Our regulon analysis predicted Smarca4 regulon activation in mutant ventricular cardiomyocytes but not in control cardiomyoctes (**Fig 2h**). Smarca4 regulon genes were enriched for signaling pathways/biologic processes such as MAPK signaling, IGF1 signaling, mTOR signaling, FAK signaling, ubiquitin mediated proteolysis, protein processing in ER, lysosome (**Fig 2k,l**). Taken together, our data in 2 HCM mouse models, and human HCM heart data suggest that the SWI/SNF complex is a therapeutic target for preventing cardiac remodeling in early-stage HCM.

### Fibrosis

Interstitial fibrosis is common in HCM^4, 56, 57^ and increases risk for heart failure, ventricular arrhythmias and sudden cardiac death^58^. Our histology results in R92W-TnT hearts, and a clinical study showing biomarkers of increased collagen synthesis in pathogenic variant carriers^59^ indicate that fibrosis can predate LVH. But the transcriptional mechanisms whereby expression of mutant sarcomeric proteins in cardiomyocytes leads to fibroblast activation and interstitial fibrosis are unclear.

Under normal physiologic conditions, fibroblasts comprise 15-24% of heart cells, and provide structural support by regulating the synthesis and degradation of ECM proteins^57^. Following TGFβ stimulation, fibroblasts proliferate and differentiate into myofibroblasts that express alpha smooth muscle actin (α-SMA) and increase expression of fibrillar collagens I/III, periostin and PDGFRa^60^. Prior studies in mouse hearts^61^ suggest activation of the TGFβ signaling pathway is important in generation of cardiac fibrosis in HCM. In our study, IPA predicted activation of fibrosis and FAK signaling only in mutant-TnT fibroblasts, which also demonstrated modest upregulation of several myofibroblast marker genes (**Suppl. Fig 4a**).

To expand our understanding of cardiac fibroblast activation and early onset of interstitial fibrosis in TnT-mutant hearts, we examined expression of genes and TFs involved in the pro-fibrotic TGFβ and Wnt signaling pathways in C22 fibroblasts which are likely to be the main effectors of interstitial fibrosis in mutant-TnT hearts, by virtue of being the largest fibroblast cluster. Mutant-TnT C22 fibroblasts demonstrated upregulation of incoming growth factor signaling (IGF, EGF, FGF, PDGF) and increased expression of several genes/TFs involved in the pro-fibrotic TGFβ and Wnt signaling pathways, but TGFβ gene (*Tgfb1-3*) expression was unchanged (**Fig 3c, d**, **Suppl. Fig 9d**). Our Cellchat analysis predicted strong activation of profibrotic periostin and TGFβ signaling only in mutant-TnT C22 fibroblasts; the source of TGFβ signaling was predicted to be C21-fibroblasts, C17-ECs, C8-cardiomyocytes, C10-epicardial cells and C1-leukocytes (**Fig 7b, c**). In contrast, in mutant-MyHC hearts, activation of TGFβ signaling was only predicted in C2-macrophages, with C11-epicardial cells as the source – this could explain lack of interstitial fibrosis in mutant-MyHC hearts at early-disease stage.

TGFβ is known to be secreted in its latent form and stored in ECM^62^. Stimulation of canonical TGFβ-SMAD signaling in fibroblasts requires activation of TGFβ prior to binding to dimeric TGFβ receptors. Prior studies suggest that, activate FAK, and canonical TGFβ-SMAD signaling which in turn can stimulate Wnt-β catenin signaling^63^ (**Suppl. Fig 11**). Abnormal cardiac mechanics (diastolic dysfunction, higher LVEF) in mutant-TnT but not mutant-MyHC at early disease stage could contribute to activation of pro-fibrotic signaling in mutant-TnT hearts, by stimulating release of active TGFβ ligand from ECM, which then binds TGFβ receptors in C22 fibroblasts, to promote collagen synthesis and interstitial fibrosis by activation of canonical TGFβ-signaling (via SMADs)^64^ and Wnt-βCatenin signaling^65^ (**Suppl. Fig 11**). Our data suggests that restoration of cardiac biomechanics (e,g, with cardiac myosin ATPase inhibitors^66^) could be antifibrotic in HCM patients with hyperdynamic LV systolic function, diastolic dysfunction and early activation of pro-fibrotic signaling pathways.

### Conclusions

Bioinformatics analysis of linked single nucleus RNA-/ATAC-seq data in two HCM mouse models predicts that ventricular myocyte hypertrophy is driven by activation of AP1 transcription factors, chromatin remodeler complexes (SWI/SNF, ISWI) and growth factor signaling pathways. Genotype-specific effects of mutant sarcomeric proteins on myofilament Ca^2+^sensitivity and cardiac mechanics, as well as intercellular communications from non-myocyte cells underlies phenotypic heterogeneity in HCM.

## MATERIALS and METHODS

All procedures involving the handling and care of mice were approved by the Animal Care and Use Committees of the University of California San Francisco, and adhered to the National Institutes of Health Public Health Service guidelines.

### Transgenic Mouse Models

The R92W-TnT^+/-^ male mouse breeders were kindly provided by Dr. Jill Tardiff (University of Arizona), and the R403Q-αMyHC^+/-^ male mouse breeders were kindly provided by Dr. Leslie Leinwand (University of Colorado Boulder). The R92W-TnT mouse is an F1 cross between FVB/N and C57/Bl6 strains^17^, and the R403Q-αMyHC were generated in CBA/B16 (F1 cross) mice^18^. The R92W-TnT^+/-^ and R403Q-MyHC^+/-^ mice were backcrossed to C57BL/6N (Charles River) for >10 generations. Male mice were weaned and genotyped at the age of 4 weeks by PCR-amplified tail DNA as described previously^15^. All studies were conducted at 5 weeks of age.

### Histology

Hearts were harvested, extracardiac vessels were trimmed and incubated in 4% PFA for 2 hours at room temperature, followed by 30% sucrose solution overnight. Subsequently, hearts were embedded in OCT and frozen at −80°C. Frozen hearts were sliced using a cryotome to generate slices of 10 µm thickness, and mounted onto glass slides. Slides were kept at −20°C until staining. Picrosirius red staining was used for quantification of fibrosis (cardiac collagen). Sections were stained with solution I (0.01% fast green FCF (Sigma, F7252) in saturated picric acid) for 1 hour, followed by solution II (0.04% fast green FCF/0.1% Sirius red (Sigma, Direct Red 80: 365548) in saturated picric acid) for 1 hour. Subsequently, sections were washed with acidified water (0.5% acetic acid), dehydrated with 100% ethanol, cleared with xylene, and mounted with DPX permanent mounting media. Stained sections were imaged under bright-field microscopy using a Leica DM6 microscope at 10x magnification. Fibrosis was quantified using ImageJ 1.53t (https://imagej.net/ij/docs/examples/stained-sections/index.html).

### Echocardiography

Cardiac phenotyping was performed by echocardiography, using a Vevo 3100 platform and MX550D 40Mhz probe (VisualSonics, Toronto, Canada). Five-week-old R92W-TnT and R403Q-MyHC mice and littermate controls were anesthetized using inhaled isoflurane (3 v/v% for induction and 0.5-1% v/v% for maintenance). Parasternal long (PLSAX) images were recorded in B-mode and M-mode. To evaluate diastolic function, pulsed-wave (PW) Doppler of mitral valve (MV) inflow, tissue Doppler imaging and left atrial (LA) area were recorded using the apical 4-chamber view. Heart rate was maintained at >450 bpm during recording of systolic parameters, and >400 bpm during recording of parameters reflecting diastolic function. Data was analyzed using the Vevo Image Lab Software (Version 5.6.0, VisualSonics, Toronto, Canada). Echocardiographic measurements were averaged from at least 3 separate cardiac cycles. Statistical Analysis was carried out using the R (version 4.1.0) package tidyverse (2.0.0) gtsummary (version 1.4.1).

### Single nucleus RNA-seq and ATAC-seq

#### Nuclei isolation

Five-week-old R92W-TnT and T403Q-MyHC mutant mice and littermate controls were imaged by echocardiography 2-4 days prior to nuclei isolation. Mutant and littermate control mice (n=4) were anaesthetized with isoflurane (3%), and euthanized by cervical dislocation. The chest cavity was opened and the heart was quickly excised along with connecting vessels, and washed in ice-cold PBS. The right and left atrial appendages, right and left ventricles were excised, cut into 1 mm^3^-sized pieces in ice-cold PBS, transferred to a 50ml conical tube containing ice-cold PBS, and centrifuged at 500g for 2 minutes at 4°C. Heart tissue pieces were resuspended in 2 ml of Lysis Buffer (Sucrose 1M, Tri-HCl pH=8 10mM, Magnesium Acetate 5mM, DTT 1mM, Triton-X 0.2%, Halt Protease Inhibitors Cocktail 1x, Rnase OUT Recombinant Ribonuclease Inhibitor 200U/ml, Nuclease-free H_2_O) in a 7.5 ml Dounce homogenizer (on ice), and homogenized to release nuclei from heart tissue. The lysate was filtered and centrifuged at 1000G for 8 minutes at 4°C. Supernatant was discarded and the pellet was resuspended in 1ml Nucleic-Buffer (Sucrose 440mM, Tri-HCl pH=7.2 10mM, Potassium Chloride 70mM, Magnesium Chloride 5mM, Spermidine 1,5mM, Halt Protease Inhibitors Cocktail 1x, Rnase OUT Recombinant Ribonuclease Inhibitor 200U/ml, Nuclease-free H_2_O), filtered and centrifuged at 1000G for 5 minutes at 4°C. The pellet was resuspended in 10XGenomics 1x Nuclear-Buffer for subsequent steps. A final count with a target concentration of 3000-8000 Nuclei/ml was obtained for library preparation, which was performed using the 10X Genomics Multiome Protocol (**Fig 1B**). Library preparation of all samples was performed in parallel. A total 50,179 nuclei passed quality control using the Cellranger Arc pipeline across all 4 experimental conditions. All four samples were sequenced together.

#### Sequencing and Postprocessing

Sequencing of ATAC and RNA-seq libraries was performed on an S4 lane of a NovaSeq 6000 sequencer, using the 10X Genomics Multiome Protocol. Fastq files were processed using Cell Ranger ARC. Cell Ranger ARC count was run on a local cluster using an MM10 reference genome. Preliminary quality control, assessment of cell number, number of high-quality fragments and genes per cell was determined by Cell Ranger ARC. The filtered feature and ATAC fragment files were used for subsequent analysis (**Fig 1B**).

#### Data Analysis

All Multiome datasets were loaded and integrated into one project using ArchR. ArchR Doublet scoring was run on the unfiltered dataset and determined doublets were removed. LSI-Dimensionality-Reduction was carried out for the ATAC and gene expression datasets separately. Subsequently ATAC and GENEX dimensions were integrated, Harmony was run on the combined dimension, and grouped by sample. UMAP projections were added for all dimensionality reductions with a minimal distance of 0.8. Clusters were determined using the Harmony dimensionality reduction, with a resolution of 1.2. Clusters were limited to 25 by Seurat-based ArchR Clustering. Pseudo bulking for subsequent analysis was performed using the AddCoverages method. Clustering and Embeddings were exported to Seurat for subsequent analysis.

#### Cluster Identity

Established cell markers, gene expression and gene score (derived from chromatin-accessibility) were employed to identify cell populations in each cluster. The marker genes used are as follows: cardiomyocytes (*Mhrt, Tnnt2, Myh6, ANP, BNP*), fibroblasts (*Col5a1, Tcf21, Pdgfra, Col3a1*), endothelial cells (*PECAM1*), vascular smooth muscle cells (*Mylk, Rgs5*), epicardial cells (Msln, Upk3b, *Gpm6a, Upk1b(7,8)*), endocardial cells (*Vwf, Igf2, H19*), macrophages (*Vsig4, C1qa*), monocytes (*CD14, CEBPB*), B cells (*Pax5, MS4A1*), T cells (*CD8A, CCL5, CD3D, Tbx21, IL7R*). Next, we examined differential gene expression in each cluster. *Gene expression analysis*: All separate datasets were integrated using the Seurat (4.0.6) Merge function, normalized and scaled after merging. Differential gene expression was determined using FindMarkers with the DESeq algorithm comparing the mutant groups to WT. Differential gene expression of both mutant groups was compared using GOPlot (1.0.2) and Venn diagrams were generated. Cell lysis leads to release of mRNA that contaminate the nuclei preparation; these mRNA can be included in the gel-bead, barcoded and sequenced. To address mRNA contamination, raw gene expression reads were analyzed using the Soup-X package (1.5.2)^67^. A contamination fraction of rho = 0.2 was set across all samples to account for differences, and prevent introduction of bias. Subsequent analysis was done using a normalized count matrix filtered using Soup-X. Soup-X determines the contamination fraction based on clustering provided by the Cell Ranger ARC pipeline; genes that were identified in all clusters are considered contaminants – these reads are subtracted across all cells.

#### Ingenuity pathway analysis (IPA)

Differential Gene expression data for both mutants and the relevant clusters was uploaded to the Qiagen IPA system (version 70750971 (Release Date: 2021-10-22), QIAGEN Inc., https://digitalinsights.qiagen.com/IPA). The mouse genome was used as a reference. All cell types and data sources that were not related to the heart were excluded, and only genes with FDR p<0.05 were considered in the downstream analysis. Activated pathways were screened and bar plots showing the 10 most up-and down-regulated pathways are shown. For canonical pathway analysis, −logP >1.3 was used as the cutoff; a z-score > or < 2 was used as the cutoff for significant pathway activation or inhibition.

#### ATAC analysis

Additional analysis of ATAC data was performed using ArchR (1.0.1)^68^ as described (10). Based on the calculated gene score, marker features were obtained and differential marker feature analysis was performed to determine differential accessibility between both mutants and littermate controls (WT). Gene tracks for multiple genes were obtained. Reproducible peaks were identified using MACS2, and a peak matrix was established. After establishing a robust peak set, we sought to predict which transcription factors (TF) drive changes in gene expression in each mutant. Motif annotations from the cisdb database were added to the peak set in ArchR, and differential testing between the mutants and WT was carried out using chromVAR to find enriched TF motifs in mutants. Only motifs with FDR p< 0.01 were considered for final analysis.

#### SCENIC analysis and cross-validation of identified transcription factor binding motifs

To evaluate drivers of gene expression in a single cell dataset, the SCENIC pipeline^69^ was developed to identify Regulons (defined as genes co-expressed with a TF factor that drives their expression). The SCENIC pipeline consists of multiple steps, based on previously published tools^69^. SCENIC uses gene expression data of a single-cell dataset to infer gene regulatory networks. In a first step using Genie3, a machine-learning based approach, co-expression of TFs and their respective putative candidate targets is predicted. Based on this data, RcisTarget is used to identify whether the potential regulators’ TF DNA-binding motif is significantly over- represented 500bp upstream of the transcription start site (TSS) of the genes of interest. Thus, regulons are built by integrating data from both steps (TF with predicted target genes). Using AUCell the gene set ‘activity’ in each cell is calculated through enrichment of the regulon as an area under the curve (AUC) across all genes in a cell based on their ranked expression value. Based on this the relative regulon activity is calculated for each cell group. Regulon activity per cell group is compared to the other cell groups **(Suppl Fig 12**). For each cluster both mutants and controls are compared, and differentially-activated regulons are determined. To validate putative driving TFs, enriched TF motifs from the ATAC data obtained in the ArchR workflow were cross-referenced with activated regulons (**Fig 1B**). Z-Score for both datasets were calculated, and a combined Z-Score (Mean of both z-scores) was generated to identify matching TFs - this cross-validation was used to identify TFs that drive cell phenotype.

#### CellChat Analysis

While analyzing differences between groups for an individual cell type renders useful information, cell-cell-interactions are often much more difficult to study. Single nucleus datasets derived from a single heart permit investigation of interactions between several different cell populations, without cell dissociation. CellChat is an R package that uses predefined clustering of the single-cell/nuclei datasets to infer cell-cell-communication using ligand-receptor-pairing based on gene expression in all cells^70^. A database with ligand-receptor interaction and their heteromeric complexes has been established and can be leveraged to predict interaction strength, pathways and signaling connections. Using laws of mass action quantification of communication probability is estimated. We used the analysis pipeline that was described in a recent publication^70^. Individual analysis was followed by differential analysis between three groups (2 mutants and littermate controls). Interactions were weighted to the number of nuclei in the respective cluster.

### Statistics

Welch’s t-test was used to compare the echocardiography and histological data of mutants with their respective littermate controls (WT); p <0.05 was considered statistically significant. Single-cell analysis was performed using Seurat, EnrichR, SCENIC for the RNAseq data and ArchR for the ATACseq data. Comparisons were made between the respective mutants and pooled littermate controls. ArchR’s getMarkerFeatures^68^ was used with Wilcoxon test. A false discovery rate (FDR) ≤ 0.05 and a Log2FC ≥ 1 were deemed relevant for differential peak accessibility, and FDR ≤ 0.01 and a MeanDiff ≥ 0.1 were deemed relevant for motif accessibility. Differential gene expression was determined in Seurat using the FindMarkers function with the DESeq2 algorithm. Genes with an adjusted p< 0.05 based on Bonferroni correction were deemed significantly dysregulated.

Using the differentially-expressed genes in the 2 mutants, we performed Ingenuity pathway analysis (IPA), or Wikipathways, or Gene Ontology (GO) term enrichment analysis to identify molecular processes that are perturbed/activated, in each cell type. For IPA canonical pathway analysis, a -log (P value) >1.3 was taken as the cutoff. A z-score >2 was defined as the cutoff for significant activation, and z-score < −2 was defined as the cutoff for significant downregulation. For pathway enrichment analysis, only DEGs with adjusted p < 0.01 were used in the analysis. Pathways with adjusted p<0.25 using the Benjamini-Hochberg method were considered for analysis. For the integrated analysis of regulon activity and motif accessibility, a combined z score was calculated from the z scores of the individual assays. For the regulon- gene pathway analysis, only matches with at least 2 overlapping genes in the respective regulon and pathways were considered. Matches were deemed significant if p < 0.01, using Fisher’s exact test.

---

**Supplemental Figure 1.** Clusteridentification. **a.** Heatmap showing the 5 top marker genes for each cluster. **b.** Bar graphs for select clusters showing the top 25 cluster-defining genes. **c.** Gene expression and chromatin accessibility w imputation of marker genes defining each cell type, projected on UMAP.

**Supplemental Figure 2.** Ventricular cardiomyocytes (C9). **a-c.** RNA-seq: Volcano plot and heat maps of top 10 up- and down-regulated DEGs in mutant TnT and MyHC cardiomyocytes normalized to controls (adj p<0.05). *Slc39a11* (Zn transmembrane transporter) and *Hbb-bs* (hemoglobin beta adult s chain) were the top 2 upregulated genes in TnT mutant cardiomyocytes (log2FC >3). **e, f.** ATAC-seq: MA plot of differentially accessible features in mutant TnT and MyHC cardiomyocytes compared to controls (FDR ≤ 0.05, log2FC ≥ 1). **g, h.** RNA-seq: Top 10 Gene Ontology biologic process enriched in up- and down-regulated genes in mutant TnT and MyHC cardiomyocytes. DNA transcription, protein phosphorylation and ubiquitin-dependent protein catabolic process were predicted to be activated in both mutant cardiomyocytes.

**Supplemental Figure 3.** C9 schematic for induction of cardiac hypertrophy in HCM. Our analysis of RNA-seq and ATAC-seq (**Fig 2c-d**, **f-g, k-i**) and echocardiography (**Fig 1d**) data leads us to hypothesize that: **a, b.** Stress activation of MAPK signaling promotes transcription of early response genes, Fos, Jun (Activator Protein 1 (AP1) transcription factors), which function as pioneer factors and recruit ATP-dependent chromatin remodeler complexes (SWI/SNF, ISWI) to nucleosome-associated enhancers - this leads to chromatin remodeling and activation of late response genes that promote myocyte hypertrophy and activation of the fetal gene program in both mutants. c. Increased intracellular Ca^2+^ activates CamKII signaling, promotes NFAT association with GATA4 and transcriptional activity of MEF2, to induce cardiac hypertrophy^99^ in both mutants. **d.** AP1 TFs promote glucocorticoid receptor binding to DNA and cardiac hypertrophy^100^ in MyHC mutants.

**Supplemental Figure 4.** Cardiac fibroblasts (Cluster C22). **a.** RNA-seq: Heat maps showing differential-expression of activated fibroblast marker genes (adj. p<0.05) in the 3 fibroblast clusters in mutant TnT and MyHC fibroblasts. Mutant TnT Cluster 22 has the highest number of DEGs, when compared to C21 and C23. Mutant MyHC fibroblasts only have 2 differentially-expressed activated fibroblast marker genes (*Col8a1, Pdgfra*) **b.** Schematic for TGFβ-mediated fibroblast activation in the 2 mutants, based on DEGs in C22 fibroblasts from both mutants. Mutant TnT fibroblasts demonstrated upregulation of several myofibroblast marker genes but not α-SMA (*Acta2*), in contrast to mutant MyHC fibroblasts that have no upregulation of myofibroblast genes. **c-f.** RNA-seq: Volcano plot and heatmap showing top 10 up- and down-regulated DEGs in mutant fibroblasts compared to controls (adj. p<0.05). *Col8a1* is the most upregulated gene in mutant TnT fibroblast cluster C22. **G-h.** ATAC-seq: Differentially accessible features (FDR ≤ 0.05, log2FC ≥ 1) in mutant TnT and MyHC fibroblasts compared to controls show stronger changes in mutant TnT. **i.** Bulk TF footprinting across mutant and control genomes show higher TF occupancy for PBX3, PGR, NR3C1 and CEBPB/D/G in mutants when compared to controls.

**Supplemental Figure 5.** Gene expression and chromatin accessibility analysis of endothelial cell cluster C14. **a-d.** RNA-seq: Volcano plots and heat maps of DEGs (adj p<0.05) for mutant TnT and MyHC ECs compared to controls **e-f.** ATAC-seq: Differentially-accessible features in mutant TnT and MyHC ECs compared to controls (FDR ≤ 0.05, Log2FC ≥ 1).

**Supplemental Figure 6.** **a.** Analysis of genes involved in cellular response to shear stress (AmiGO gene list) reveal greater dysregulation of gene expression in mutant TnT when compared to mutant MyHC ECs. **b. Schematic.** Our analysis of RNA-seq and ATAC-seq data (**Figs 4c,d**, **g-k, Suppl Fig 6a**) leads us to hypothesize that hyperdynamic LV function increases shear stress and circumferential stress in coronaries, which activates mechanosensitive receptors such as integrins, FAK signaling which leads to EGFR transactivation. This leads to activation of STAT and ERK signaling, that promote Mef2 phosphorylation and transcription of Klf2/4, which leads to activation of downstream genes such as NOS3 and increased NO production by ECs.

**Supplemental Figure 7.** Cardiac macrophages (C2). **a-d.** RNA-seq: Volcano plots and heat maps of DEGs (adj p<0.05) in mutants compared to controls. **e, f.** ATAC-seq: Differentially-accessible features ((FDR ≤ 0.05, log2FC ≥ 1) in mutant TnT and MyHC show greater changes in mutant TnT.

**Supplemental Figure 8.** Schematic. Our analysis of RNA- and ATAC-seq data in cardiac macrophages leads us to hypothesize the following. **a.** Greater activation of IRF5, REL (NFkB), IRF regulons in mutant TnT macrophages (**Fig 6h**) suggest greater macrophage polarization^101^ towards M1 in mutant TnT when compared to mutant MyHC. **b.** Damaged mitochondria and other cargo from mutant myocytes is phagocytosed by resident cardiac macrophages and degraded by phagolysosomes (**Fig 6c-d**, **k**), which promotes cardiac myocyte homeostasis^59^. **c.** Activation of pattern recognition receptors such as TLR (toll-like receptors) and NOD (nucleotide binding and oligomerization domain-like receptors) by damaged mitochondria from cardiac myocytes, leads to nuclear translocation and DNA binding of IRFs, NF-kB (REL) (**Fig 6h-i**) which is pro-inflammatory. Simultaneous activation of the glucocorticoid receptor (NR3C1) in mutant macrophages (**Fig 6h**) blocks the pro-inflammatory effects of IRF and NF-kB activation.

**Supplemental Figure 9.** Cell-Cell-Communication Analysis with CellChat (RNA-seq). **a.** Communication between clusters shown for both mutants and the combined controls. Colored dots represent each cluster with color of connecting line showing origin of signaling. Cell types are grouped together. Stronger and added connections in both mutants compared to controls. **b, c.** Heatmap showing predicted outgoing signaling in controls and mutants for each detected signaling pathway. The left lower quadrant shows signaling that is only activated in the mutants can be seen. d, e. Heatmap showing predicted incoming signaling in controls and mutants for each detected signaling pathway. The left lower quadrant shows signaling that is only activated in the mutants can be seen.

**Supplemental Figure 10.** RNA-seq: Comparison of predicted signaling in mutants for NPR1, VCAM, GAS, AGRN, EDN, PDGF. Clusters are represented as dots and line color depicts the originating cluster of the depicted signaling. **a.** NPR1 signaling is predicted to originate from several clusters, with C9 and C22 being the major senders; signaling is received by the epicardial cell cluster C12 and the fibroblast cluster C21 in mutant TnT, and by the epicardial cell cluster C11 in mutant MyHC. **b.** VCAM signaling is predicted to originate from several clusters, and is received by C16 in both mutants. **c.** GAS signaling is predicted to originate from C9 and C16 in the TnT mutant, and from C8 and C17 in the MyHC mutant; major signaling recipients are the macrophage cluster C2 and the fibroblast cluster C22, in both mutants. **d.** AGRN signaling originates from C8 in mutant TnT, and C9 in mutant MyHC; main recipient in mutant TnT is the C8 cardiomyocyte cluster and C9 (autocrine signaling) in mutant MyHC. **e.** EDN (endothelin) signaling is only predicted in TnT. EDN signaling originates from the fibroblast cluster C21, and is received by the ventricular cardiomyocyte cluster C9. **f.** PDGF signaling mainly originates from endothelial cell cluster C14 and is received by the VSMC cell cluster C20 in both mutants, and the fibroblast cluster C22 only in TnT.

**Supplemental Figure 11.** Schematic for myofibroblast activation and stimulation of interstitial fibrosis in mutant TnT hearts. Our RNA-/ATAC-seq and echocardiography data led us to hypothesize that hyperdynamic LV function/diastolic dysfunction (**Fig 1d**) promote release of active TGFß in from cardiac extracellular matrix. TGFß binds TGFBR on fibroblasts, which activates SMADs (**Fig 3d**) and MAPK signaling (**Fig 3**j). Activated SMADs translocate to the nucleus where they associate with cofactors such as EP300, CREBBP to induce expression of collagen genes^87^ (**Fig 3d**). Growth factors such as IGF, EGF, FGF, PDGF can also activate MAPK signaling (**Fig 3j**) by binding receptor tyrosine kinase (RTK) receptors on fibroblasts. MAPK activation (**Fig3d**) can stimulate Wnt-ß-catenin signaling. Beta-catenin translocates to the nucleus, where it associates with TCF4 and FOXP2 (**Fig 3d-i**) to promote transcription of cell cycle genes and Wnt target genes^88^, thus potentiating the pro-fibrotic effect of activated TGFβ signaling.

**Supplemental Figure 12.** Regulon activity across all clusters and experimental conditions. Names of regulons are given on the right with the number of regulated genes in parenthesis. Genes are shown in alphabetical order. Individual cell types are delineated with black lines.

## Supplementary information

Supplementary information is available online.

## Acknowledgments

This study was supported by the Spieker Foundation and startup funds from the UCSF division of Cardiology to MRA. Dr. Thottakara was supported by a grant from the German Heart Foundation. We would like to thank Benoit G. Bruneau, PhD for his value advise for this manuscript.

## Author Contributions

**Conceptualisation:** Ti.T., M.R.A, A.P., J.E.O., C.S.L., **Investigation:** Ti.T., M.R.A., A.P., Ta.T., Th.T., J.D., S. R. A., **Data analysis:** Ti.T. **Visualisation:** Ti.T., M.R.A., A.P., **Writing—original draft:** Ti.T., M.R.A **and writing—review and editing:** Ti.T., M.R.A., A.P., C.S.L.

## Bibliography

1. Maron BJ. Hypertrophic cardiomyopathy: a systematic review. JAMA. 2002;287:1308–20.

2. Maron BJ, Wolfson JK, Epstein SE and Roberts WC. Intramural (“small vessel”) coronary artery disease in hypertrophic cardiomyopathy. J Am Coll Cardiol. 1986;8:545–57.

3. Yalcin H, Valenta I, Yalcin F, Corona-Villalobos C, Vasquez N, Ra J, Kucukler N, Tahari A, Pozios I, Zhou Y, Pomper M, Abraham TP, Schindler TH and Abraham MR. Effect of Diffuse Subendocardial Hypoperfusion on Left Ventricular Cavity Size by (13)N-Ammonia Perfusion PET in Patients With Hypertrophic Cardiomyopathy. Am J Cardiol. 2016;118:1908–1915.

4. Sivalokanathan S, Zghaib T, Greenland GV, Vasquez N, Kudchadkar SM, Kontari E, Lu DY, Dolores-Cerna K, van der Geest RJ, Kamel IR, Olgin JE, Abraham TP, Nazarian S, Zimmerman SL and Abraham MR. Hypertrophic Cardiomyopathy Patients With Paroxysmal Atrial Fibrillation Have a High Burden of Left Atrial Fibrosis by Cardiac Magnetic Resonance Imaging. JACC Clin Electrophysiol. 2019;5:364–375.

5. Lu DY, Ventoulis I, Liu H, Kudchadkar SM, Greenland GV, Yalcin H, Kontari E, Goyal S, Corona-Villalobos CP, Vakrou S, Zimmerman SL, Abraham TP and Abraham MR. Sex-specific cardiac phenotype and clinical outcomes in patients with hypertrophic cardiomyopathy. Am Heart J. 2020;219:58–69.

6. Chaffin M, Papangeli I, Simonson B, Akkad AD, Hill MC, Arduini A, Fleming SJ, Melanson M, Hayat S, Kost-Alimova M, Atwa O, Ye J, Bedi KC, Jr., Nahrendorf M, Kaushik VK, Stegmann CM, Margulies KB, Tucker NR and Ellinor PT. Single-nucleus profiling of human dilated and hypertrophic cardiomyopathy. Nature. 2022;608:174–180.

7. Larson A, Codden CJ, Huggins GS, Rastegar H, Chen FY, Maron BJ, Rowin EJ, Maron MS and Chin MT. Altered intercellular communication and extracellular matrix signaling as a potential disease mechanism in human hypertrophic cardiomyopathy. Sci Rep. 2022;12:5211.

8. Bos JM, Hebl VB, Oberg AL, Sun Z, Herman DS, Teekakirikul P, Seidman JG, Seidman CE, Dos Remedios CG, Maleszewski JJ, Schaff HV, Dearani JA, Noseworthy PA, Friedman PA, Ommen SR, Brozovich FV and Ackerman MJ. Marked Up-Regulation of ACE2 in Hearts of Patients With Obstructive Hypertrophic Cardiomyopathy: Implications for SARS-CoV-2-Mediated COVID-19. Mayo Clin Proc. 2020;95:1354–1368.

9. Liu Y, Afzal J, Vakrou S, Greenland GV, Talbot CC, Jr., Hebl VB, Guan Y, Karmali R, Tardiff JC, Leinwand LA, Olgin JE, Das S, Fukunaga R and Abraham MR. Differences in microRNA-29 and Pro-fibrotic Gene Expression in Mouse and Human Hypertrophic Cardiomyopathy. Front Cardiovasc Med. 2019;6:170.

10. Vakrou S, Liu Y, Zhu L, Greenland GV, Simsek B, Hebl VB, Guan Y, Woldemichael K, Talbot CC, Aon MA, Fukunaga R and Abraham MR. Differences in molecular phenotype in mouse and human hypertrophic cardiomyopathy. Sci Rep. 2021;11:13163.

11. Previs MJ, O’Leary TS, Morley MP, Palmer BM, LeWinter M, Yob JM, Pagani FD, Petucci C, Kim MS, Margulies KB, Arany Z, Kelly DP and Day SM. Defects in the Proteome and Metabolome in Human Hypertrophic Cardiomyopathy. Circ Heart Fail. 2022;15:e009521.

12. Ranjbarvaziri S, Kooiker KB, Ellenberger M, Fajardo G, Zhao M, Vander Roest AS, Woldeyes RA, Koyano TT, Fong R, Ma N, Tian L, Traber GM, Chan F, Perrino J, Reddy S, Chiu W, Wu JC, Woo JY, Ruppel KM, Spudich JA, Snyder MP, Contrepois K and Bernstein D. Altered Cardiac Energetics and Mitochondrial Dysfunction in Hypertrophic Cardiomyopathy. Circulation. 2021;144:1714–1731.

13. Wang W, Wang J, Yao K, Wang S, Nie M, Zhao Y, Wang B, Pang H, Xu J, Wu G, Lu M, Tang N, Qi C, Pei H, Luo X, Li D, Yang T, Sun Q, Wei X, Li Y, Jiang D, Li P, Song L and Hu Z. Metabolic characterization of hypertrophic cardiomyopathy in human heart. Nature Cardiovascular Research. 2022;1:445–461.

14. Kawana M, Spudich JA and Ruppel KM. Hypertrophic cardiomyopathy: Mutations to mechanisms to therapies. Front Physiol. 2022;13:975076.

15. Vakrou S, Fukunaga R, Foster DB, Sorensen L, Liu Y, Guan Y, Woldemichael K, Pineda-Reyes R, Liu T, Tardiff JC, Leinwand LA, Tocchetti CG, Abraham TP, O’Rourke B, Aon MA and Abraham MR. Allele-specific differences in transcriptome, miRNome, and mitochondrial function in two hypertrophic cardiomyopathy mouse models. JCI Insight. 2018;3.

16. Ertz-Berger BR, He H, Dowell C, Factor SM, Haim TE, Nunez S, Schwartz SD, Ingwall JS and Tardiff JC. Changes in the chemical and dynamic properties of cardiac troponin T cause discrete cardiomyopathies in transgenic mice. Proc Natl Acad Sci U S A. 2005;102:18219–24.

17. Tardiff JC, Hewett TE, Palmer BM, Olsson C, Factor SM, Moore RL, Robbins J and Leinwand LA. Cardiac troponin T mutations result in allele-specific phenotypes in a mouse model for hypertrophic cardiomyopathy. J Clin Invest. 1999;104:469–81.

18. Vikstrom KL, Factor SM and Leinwand LA. Mice expressing mutant myosin heavy chains are a model for familial hypertrophic cardiomyopathy. Mol Med. 1996;2:556–67.

19. Moolman JC, Corfield VA, Posen B, Ngumbela K, Seidman C, Brink PA and Watkins H. Sudden death due to troponin T mutations. J Am Coll Cardiol. 1997;29:549–55.

20. Abraham MR, Bottomley PA, Dimaano VL, Pinheiro A, Steinberg A, Traill TA, Abraham TP and Weiss RG. Creatine kinase adenosine triphosphate and phosphocreatine energy supply in a single kindred of patients with hypertrophic cardiomyopathy. Am J Cardiol. 2013;112:861–6.

21. Geisterfer-Lowrance AA, Kass S, Tanigawa G, Vosberg HP, McKenna W, Seidman CE and Seidman JG. A molecular basis for familial hypertrophic cardiomyopathy: a beta cardiac myosin heavy chain gene missense mutation. Cell. 1990;62:999–1006.

22. Becker KD, Gottshall KR, Hickey R, Perriard JC and Chien KR. Point mutations in human beta cardiac myosin heavy chain have differential effects on sarcomeric structure and assembly: an ATP binding site change disrupts both thick and thin filaments, whereas hypertrophic cardiomyopathy mutations display normal assembly. J Cell Biol. 1997;137:131–40.

23. Volkmann N, Lui H, Hazelwood L, Trybus KM, Lowey S and Hanein D. The R403Q myosin mutation implicated in familial hypertrophic cardiomyopathy causes disorder at the actomyosin interface. PLoS One. 2007;2:e1123.

24. Colegrave M and Peckham M. Structural implications of beta-cardiac myosin heavy chain mutations in human disease. Anat Rec (Hoboken). 2014;297:1670–80.

25. Lowey S, Bretton V, Joel PB, Trybus KM, Gulick J, Robbins J, Kalganov A, Cornachione AS and Rassier DE. Hypertrophic cardiomyopathy R403Q mutation in rabbit beta-myosin reduces contractile function at the molecular and myofibrillar levels. Proc Natl Acad Sci U S A. 2018;115:11238–11243.

26. Fatkin D, McConnell BK, Mudd JO, Semsarian C, Moskowitz IG, Schoen FJ, Giewat M, Seidman CE and Seidman JG. An abnormal Ca(2+) response in mutant sarcomere protein-mediated familial hypertrophic cardiomyopathy. J Clin Invest. 2000;106:1351–9.

27. Geisterfer-Lowrance AA, Christe M, Conner DA, Ingwall JS, Schoen FJ, Seidman CE and Seidman JG. A mouse model of familial hypertrophic cardiomyopathy. Science. 1996;272:731–4.

28. Garces de Los Fayos Alonso I, Liang HC, Turner SD, Lagger S, Merkel O and Kenner L. The Role of Activator Protein-1 (AP-1) Family Members in CD30-Positive Lymphomas. Cancers (Basel). 2018;10.

29. Bevilacqua A, Willis MS and Bultman SJ. SWI/SNF chromatin-remodeling complexes in cardiovascular development and disease. Cardiovasc Pathol. 2014;23:85–91.

30. Reyes AA, Marcum RD and He Y. Structure and Function of Chromatin Remodelers. J Mol Biol. 2021;433:166929.

31. Kreusser MM and Backs J. Integrated mechanisms of CaMKII-dependent ventricular remodeling. Front Pharmacol. 2014;5:36.

32. Sampieri L, Di Giusto P and Alvarez C. CREB3 Transcription Factors: ER-Golgi Stress Transducers as Hubs for Cellular Homeostasis. Front Cell Dev Biol. 2019;7:123.

33. Groenendijk BC, Van der Heiden K, Hierck BP and Poelmann RE. The role of shear stress on ET-1, KLF2, and NOS-3 expression in the developing cardiovascular system of chicken embryos in a venous ligation model. Physiology (Bethesda). 2007;22:380–9.

34. Qi YX, Han Y and Jiang ZL. Mechanobiology and Vascular Remodeling: From Membrane to Nucleus. Adv Exp Med Biol. 2018;1097:69–82.

35. Shyy JY and Chien S. Role of integrins in endothelial mechanosensing of shear stress. Circ Res. 2002;91:769–75.

36. Makki N, Thiel KW and Miller FJ, Jr. The epidermal growth factor receptor and its ligands in cardiovascular disease. Int J Mol Sci. 2013;14:20597–613.

37. Andersen P, Pedersen MW, Woetmann A, Villingshoj M, Stockhausen MT, Odum N and Poulsen HS. EGFR induces expression of IRF-1 via STAT1 and STAT3 activation leading to growth arrest of human cancer cells. Int J Cancer. 2008;122:342–9.

38. Atkins GB and Jain MK. Role of Kruppel-like transcription factors in endothelial biology. Circ Res. 2007;100:1686–95.

39. Lee JH, Chun T, Park SY and Rho SB. Interferon regulatory factor-1 (IRF-1) regulates VEGF-induced angiogenesis in HUVECs. Biochim Biophys Acta. 2008;1783:1654–62.

40. Nicolas-Avila JA, Lechuga-Vieco AV, Esteban-Martinez L, Sanchez-Diaz M, Diaz-Garcia E, Santiago DJ, Rubio-Ponce A, Li JL, Balachander A, Quintana JA, Martinez-de-Mena R, Castejon-Vega B, Pun-Garcia A, Traves PG, Bonzon-Kulichenko E, Garcia-Marques F, Cusso L, N AG, Gonzalez-Guerra A, Roche-Molina M, Martin-Salamanca S, Crainiciuc G, Guzman G, Larrazabal J, Herrero-Galan E, Alegre-Cebollada J, Lemke G, Rothlin CV, Jimenez-Borreguero LJ, Reyes G, Castrillo A, Desco M, Munoz-Canoves P, Ibanez B, Torres M, Ng LG, Priori SG, Bueno H, Vazquez J, Cordero MD, Bernal JA, Enriquez JA and Hidalgo A. A Network of Macrophages Supports Mitochondrial Homeostasis in the Heart. Cell. 2020;183:94–109 e23.

41. Gong T, Liu L, Jiang W and Zhou R. DAMP-sensing receptors in sterile inflammation and inflammatory diseases. Nat Rev Immunol. 2020;20:95–112.

42. Platanitis E and Decker T. Regulatory Networks Involving STATs, IRFs, and NFkappaB in Inflammation. Front Immunol. 2018;9:2542.

43. Karsenty C, Guilbeau-Frugier C, Genet G, Seguelas MH, Alzieu P, Cazorla O, Montagner A, Blum Y, Dubroca C, Maupoint J, Tramunt B, Cauquil M, Sulpice T, Richard S, Arcucci S, Flores-Flores R, Pataluch N, Montoriol R, Sicard P, Deney A, Couffinhal T, Senard JM and Gales C. Ephrin-B1 regulates the adult diastolic function through a late postnatal maturation of cardiomyocyte surface crests. Elife. 2023;12.

44. Zurek M, Johansson E, Palmer M, Albery T, Johansson K, Ryden-Markinhutha K and Wang QD. Neuregulin-1 Induces Cardiac Hypertrophy and Impairs Cardiac Performance in Post-Myocardial Infarction Rats. Circulation. 2020;142:1308–1311.

45. Manivasagam S and Vellaichamy E. Suppression of Npr1, not Npr2 gene function induces hypertrophic growth in H9c2 cells in vitro. Biochem Biophys Res Commun. 2017;491:250–256.

46. Lu DY, Pozios I, Haileselassie B, Ventoulis I, Liu H, Sorensen LL, Canepa M, Phillip S, Abraham MR and Abraham TP. Clinical Outcomes in Patients With Nonobstructive, Labile, and Obstructive Hypertrophic Cardiomyopathy. J Am Heart Assoc. 2018;7.

47. Buttner P, Ueberham L, Shoemaker MB, Roden DM, Dinov B, Hindricks G, Bollmann A and Husser D. Identification of Central Regulators of Calcium Signaling and ECM-Receptor Interaction Genetically Associated With the Progression and Recurrence of Atrial Fibrillation. Front Genet. 2018;9:162.

48. Semsarian C, Ahmad I, Giewat M, Georgakopoulos D, Schmitt JP, McConnell BK, Reiken S, Mende U, Marks AR, Kass DA, Seidman CE and Seidman JG. The L-type calcium channel inhibitor diltiazem prevents cardiomyopathy in a mouse model. J Clin Invest. 2002;109:1013–20.

49. Kataoka A, Hemmer C and Chase PB. Computational simulation of hypertrophic cardiomyopathy mutations in troponin I: influence of increased myofilament calcium sensitivity on isometric force, ATPase and [Ca2+]i. J Biomech. 2007;40:2044–52.

50. Guinto PJ, Haim TE, Dowell-Martino CC, Sibinga N and Tardiff JC. Temporal and mutation-specific alterations in Ca2+ homeostasis differentially determine the progression of cTnT-related cardiomyopathies in murine models. Am J Physiol Heart Circ Physiol. 2009;297:H614–26.

51. Mehrotra A, Joe B and de la Serna IL. SWI/SNF chromatin remodeling enzymes are associated with cardiac hypertrophy in a genetic rat model of hypertension. J Cell Physiol. 2013;228:2337–42.

52. Taimor G, Schluter KD, Best P, Helmig S and Piper HM. Transcription activator protein 1 mediates alpha- but not beta-adrenergic hypertrophic growth responses in adult cardiomyocytes. Am J Physiol Heart Circ Physiol. 2004;286:H2369–75.

53. Otto JE, Ursu O, Wu AP, Winter EB, Cuoco MS, Ma S, Qian K, Michel BC, Buenrostro JD, Berger B, Regev A and Kadoch C. Structural and functional properties of mSWI/SNF chromatin remodeling complexes revealed through single-cell perturbation screens. Mol Cell. 2023;83:1350–1367 e7.

54. Hang CT, Yang J, Han P, Cheng HL, Shang C, Ashley E, Zhou B and Chang CP. Chromatin regulation by Brg1 underlies heart muscle development and disease. Nature. 2010;466:62–7.

55. Scherba JC, Halushka MK, Andersen ND, Maleszewski JJ, Landstrom AP, Bursac N and Glass C. BRG1 is a biomarker of hypertrophic cardiomyopathy in human heart specimens. Sci Rep. 2022;12:7996.

56. Hurtado-de-Mendoza D, Corona-Villalobos CP, Pozios I, Gonzales J, Soleimanifard Y, Sivalokanathan S, Montoya-Cerrillo D, Vakrou S, Kamel I, Mormontoy-Laurel W, Dolores-Cerna K, Suarez J, Perez-Melo S, Bluemke DA, Abraham TP, Zimmerman SL and Abraham MR. Diffuse interstitial fibrosis assessed by cardiac magnetic resonance is associated with dispersion of ventricular repolarization in patients with hypertrophic cardiomyopathy. J Arrhythm. 2017;33:201–207.

57. Schlittler M, Pramstaller PP, Rossini A and De Bortoli M. Myocardial Fibrosis in Hypertrophic Cardiomyopathy: A Perspective from Fibroblasts. Int J Mol Sci. 2023;24.

58. Shirani J, Pick R, Roberts WC and Maron BJ. Morphology and significance of the left ventricular collagen network in young patients with hypertrophic cardiomyopathy and sudden cardiac death. J Am Coll Cardiol. 2000;35:36–44.

59. Ho CY, Lopez B, Coelho-Filho OR, Lakdawala NK, Cirino AL, Jarolim P, Kwong R, Gonzalez A, Colan SD, Seidman JG, Diez J and Seidman CE. Myocardial fibrosis as an early manifestation of hypertrophic cardiomyopathy. N Engl J Med. 2010;363:552–63.

60. Gibb AA, Lazaropoulos MP and Elrod JW. Myofibroblasts and Fibrosis: Mitochondrial and Metabolic Control of Cellular Differentiation. Circ Res. 2020;127:427–447.

61. Teekakirikul P, Eminaga S, Toka O, Alcalai R, Wang L, Wakimoto H, Nayor M, Konno T, Gorham JM, Wolf CM, Kim JB, Schmitt JP, Molkentin JD, Norris RA, Tager AM, Hoffman SR, Markwald RR, Seidman CE and Seidman JG. Cardiac fibrosis in mice with hypertrophic cardiomyopathy is mediated by non-myocyte proliferation and requires Tgf-beta. J Clin Invest. 2010;120:3520–9.

62. Shi M, Zhu J, Wang R, Chen X, Mi L, Walz T and Springer TA. Latent TGF-beta structure and activation. Nature. 2011;474:343–9.

63. Dzialo E, Tkacz K and Blyszczuk P. Crosstalk between the TGF-beta and WNT signalling pathways during cardiac fibrogenesis. Acta Biochim Pol. 2018;65:341–349.

64. Khalil H, Kanisicak O, Prasad V, Correll RN, Fu X, Schips T, Vagnozzi RJ, Liu R, Huynh T, Lee SJ, Karch J and Molkentin JD. Fibroblast-specific TGF-beta-Smad2/3 signaling underlies cardiac fibrosis. J Clin Invest. 2017;127:3770–3783.

65. Richter G, Gui T, Bourgeois B, Koyani CN, Ulz P, Heitzer E, von Lewinski D, Burgering BMT, Malle E and Madl T. beta-catenin regulates FOXP2 transcriptional activity via multiple binding sites. FEBS J. 2021;288:3261–3284.

66. Braunwald E, Saberi S, Abraham TP, Elliott PM and Olivotto I. Mavacamten: a first-in-class myosin inhibitor for obstructive hypertrophic cardiomyopathy. Eur Heart J. 2023;44:4622–4633.

67. Young MD and Behjati S. SoupX removes ambient RNA contamination from droplet-based single-cell RNA sequencing data. Gigascience. 2020;9.

68. Granja JM, Corces MR, Pierce SE, Bagdatli ST, Choudhry H, Chang HY and Greenleaf WJ. ArchR is a scalable software package for integrative single-cell chromatin accessibility analysis. Nat Genet. 2021;53:403–411.

69. Aibar S, Gonzalez-Blas CB, Moerman T, Huynh-Thu VA, Imrichova H, Hulselmans G, Rambow F, Marine JC, Geurts P, Aerts J, van den Oord J, Atak ZK, Wouters J and Aerts S. SCENIC: single-cell regulatory network inference and clustering. Nat Methods. 2017;14:1083–1086.

70. Jin S, Guerrero-Juarez CF, Zhang L, Chang I, Ramos R, Kuan CH, Myung P, Plikus MV and Nie Q. Inference and analysis of cell-cell communication using CellChat. Nat Commun. 2021;12:1088.

