## Supplementary Figures for "Single-nucleus RNA/ATAC-seq in early-stage HCM models predicts SWI/SNF-activation in mutant-myocytes, and allele-specific differences in fibroblasts"

#### Suppl. Fig 1

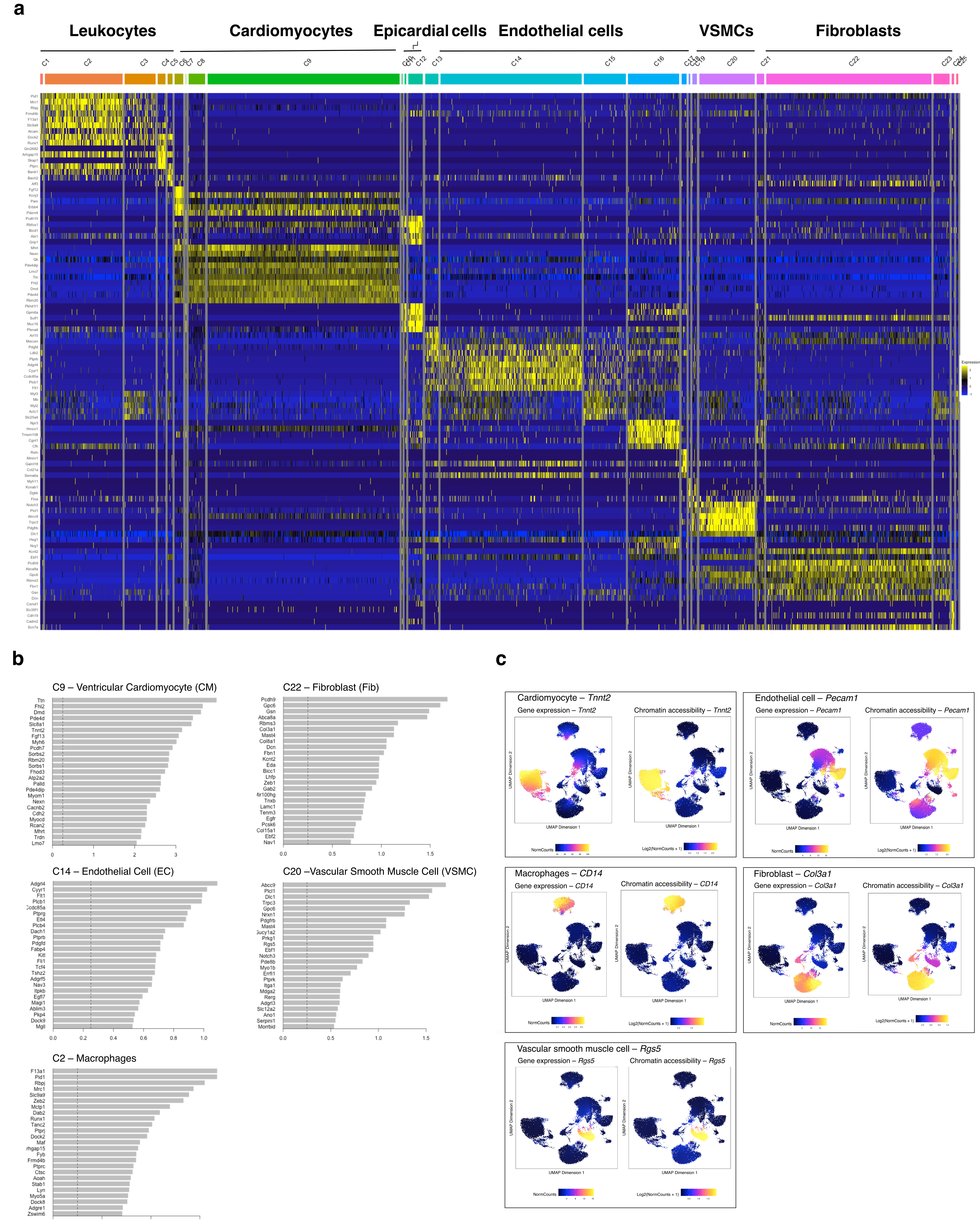


**Supplemental Figure 1**

**a.** Heatmap showing the 5 top marker genes for each cluster. **b.** Bar graphs for select clusters showing the top 25 cluster-defining genes. **c.** Gene expression and chromatin accessibility w imputation of marker genes defining each cell type, projected on UMAP.

**Suppl. Fig 2**

### C9 - Ventricular Cardiomyocytes

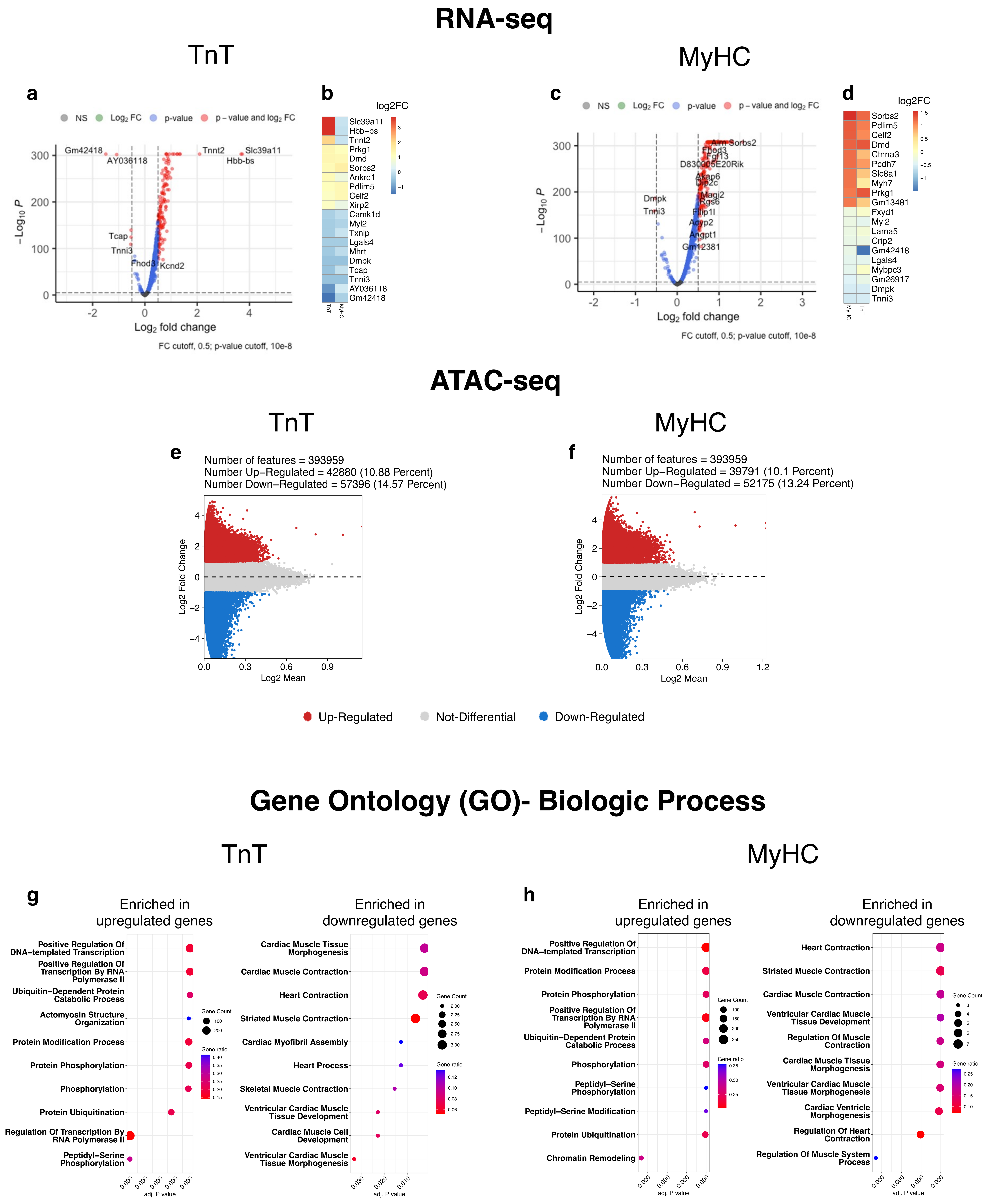


**Supplemental Figure 2. Ventricular cardiomyocytes (C9).**

**a-c.** RNA-seq: Volcano plot and heat maps of top 10 up- and down-regulated DEGs in mutant TnT and MyHC cardiomyocytes normalized to controls (adj $p<0.05$). *Slc39a11* (Zn transmembrane transporter) and *Hbb-bs* (hemoglobin beta adult s chain) were the top 2 upregulated genes in TnT mutant cardiomyocytes (log2FC >3). **e, f.** ATAC-seq: MA plot of differentially accessible features in mutant TnT and MyHC cardiomyocytes compared to controls (FDR ≤ 0.05, log2FC ≥ 1). **g, h.** RNA-seq: Top 10 Gene Ontology biologic process enriched in up- and down-regulated genes in mutant TnT and MyHC cardiomyocytes. DNA transcription, protein phosphorylation and ubiquitin-dependent protein catabolic process were predicted to be activated in both mutant cardiomyocytes.

Suppl. Fig 3

#### Ventricular Cardiomyocytes Mechanism

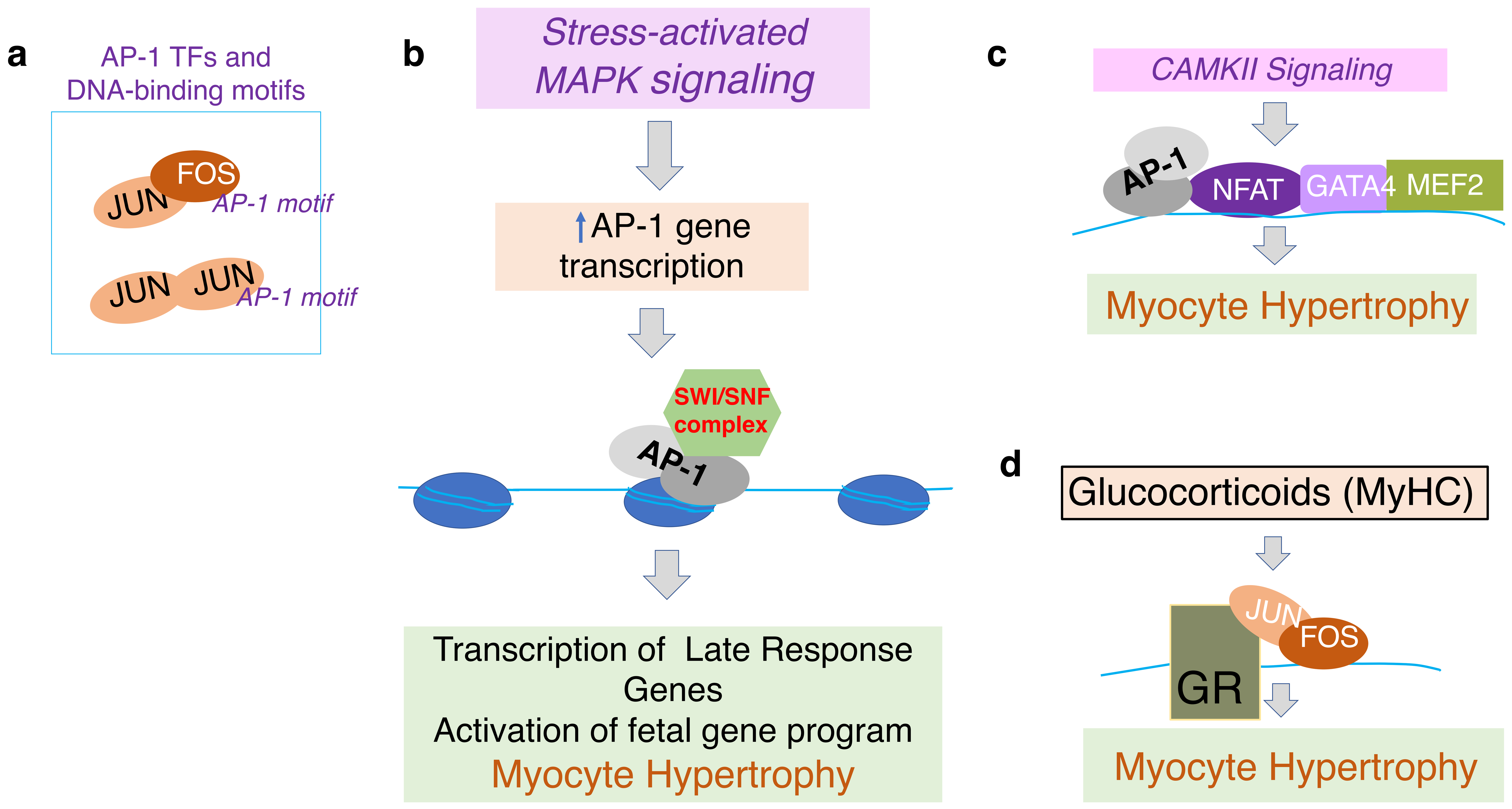


**Supplemental Figure 3. C9 schematic for induction of cardiac hypertrophy in HCM.**

Our analysis of RNA-seq and ATAC-seq (**Fig 2c-d, f-g, k-i**) and echocardiography (**Fig 1d**) data leads us to hypothesize that: **a, b.** Stress activation of MAPK signaling promotes transcription of early response genes, Fos, Jun (Activator Protein 1 (AP1) transcription factors), which function as pioneer factors and recruit ATP-dependent chromatin remodeler complexes (SWI/SNF, ISWI) to nucleosome-associated enhancers - this leads to chromatin remodeling and activation of late response genes that promote myocyte hypertrophy and activation of the fetal gene program in both mutants. c. Increased intracellular $Ca^{2+}$ activates CamKII signaling, promotes NFAT association with GATA4 and transcriptional activity of MEF2, to induce cardiac hypertrophy[99] in both mutants. **d.** AP1 TFs promote glucocorticoid receptor binding to DNA and cardiac hypertrophy[100] in MyHC mutants.

#### Suppl. Fig 4

### C22 - Fibroblasts

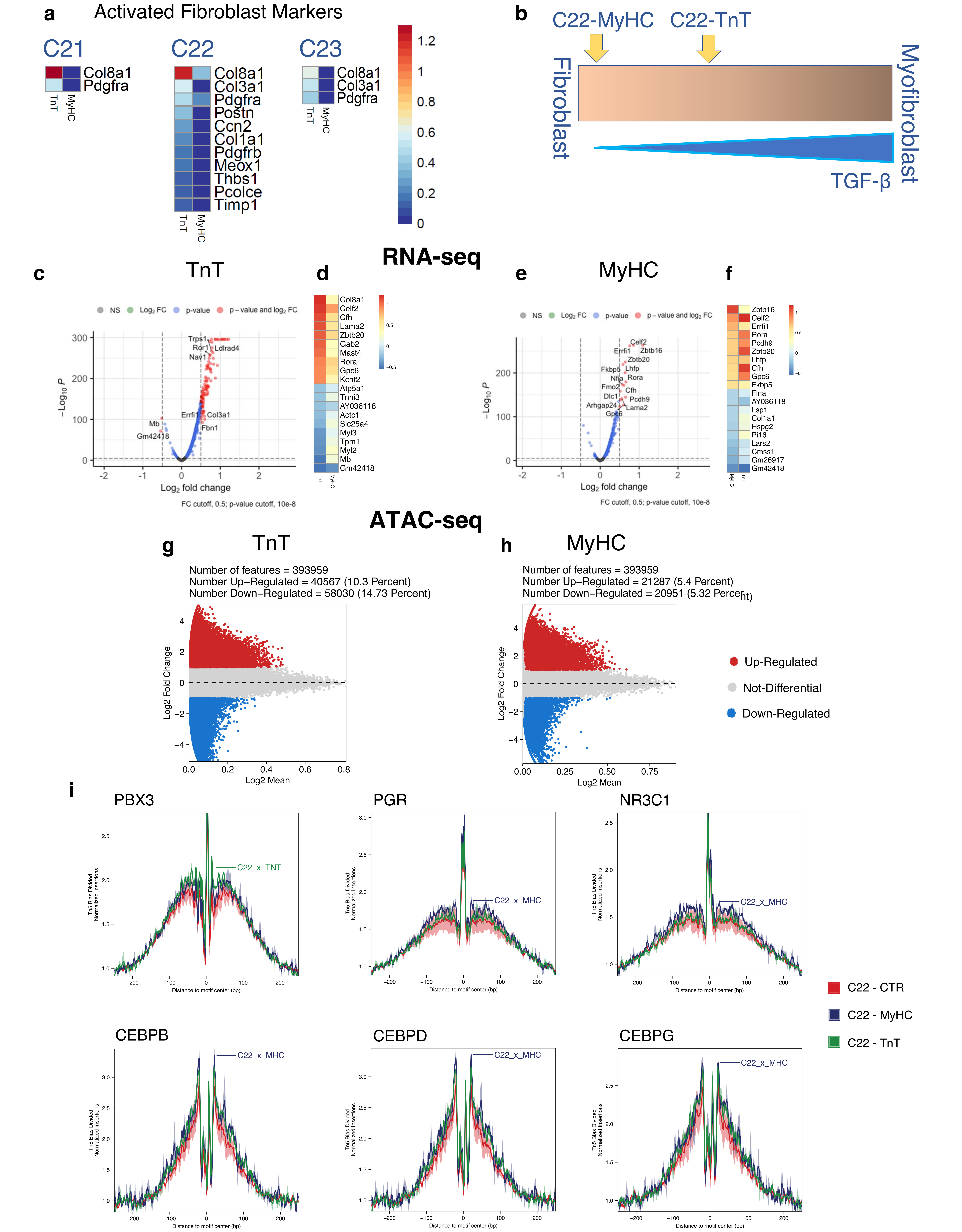


**Supplemental Figure 4. Cardiac fibroblasts (Cluster C22).**

**a.** RNA-seq: Heat maps showing differential-expression of activated fibroblast marker genes (adj. p<0.05) in the 3 fibroblast clusters in mutant TnT and MyHC fibroblasts. Mutant TnT Cluster 22 has the highest number of DEGs, when compared to C21 and C23. Mutant MyHC fibroblasts only have 2 differentially-expressed activated fibroblast marker genes (*Col8a1, Pdgfra*) **b.** Schematic for TGFβ-mediated fibroblast activation in the 2 mutants, based on DEGs in C22 fibroblasts from both mutants. Mutant TnT fibroblasts demonstrated upregulation of several myofibroblast marker genes but not α-SMA (*Acta2*), in contrast to mutant MyHC fibroblasts that have no upregulation of myofibroblast genes. **c-f.** RNA-seq: Volcano plot and heatmap showing top 10 up- and down-regulated DEGs in mutant fibroblasts compared to controls (adj. p<0.05). *Col8a1* is the most upregulated gene in mutant TnT fibroblast cluster C22. **g.**/h.ATAC-seq: Differentially accessible features (FDR ≤ 0.05, log2FC ≥ 1) in mutant TnT and MyHC fibroblasts compared to controls show stronger changes in mutant TnT. **i.** Bulk TF footprinting across mutant and control genomes show higher TF occupancy for PBX3, PGR, NR3C1 and CEBPB/D/G in mutants when compared to controls.

**Suppl. Fig 5**

### C14 - Endothelial cells

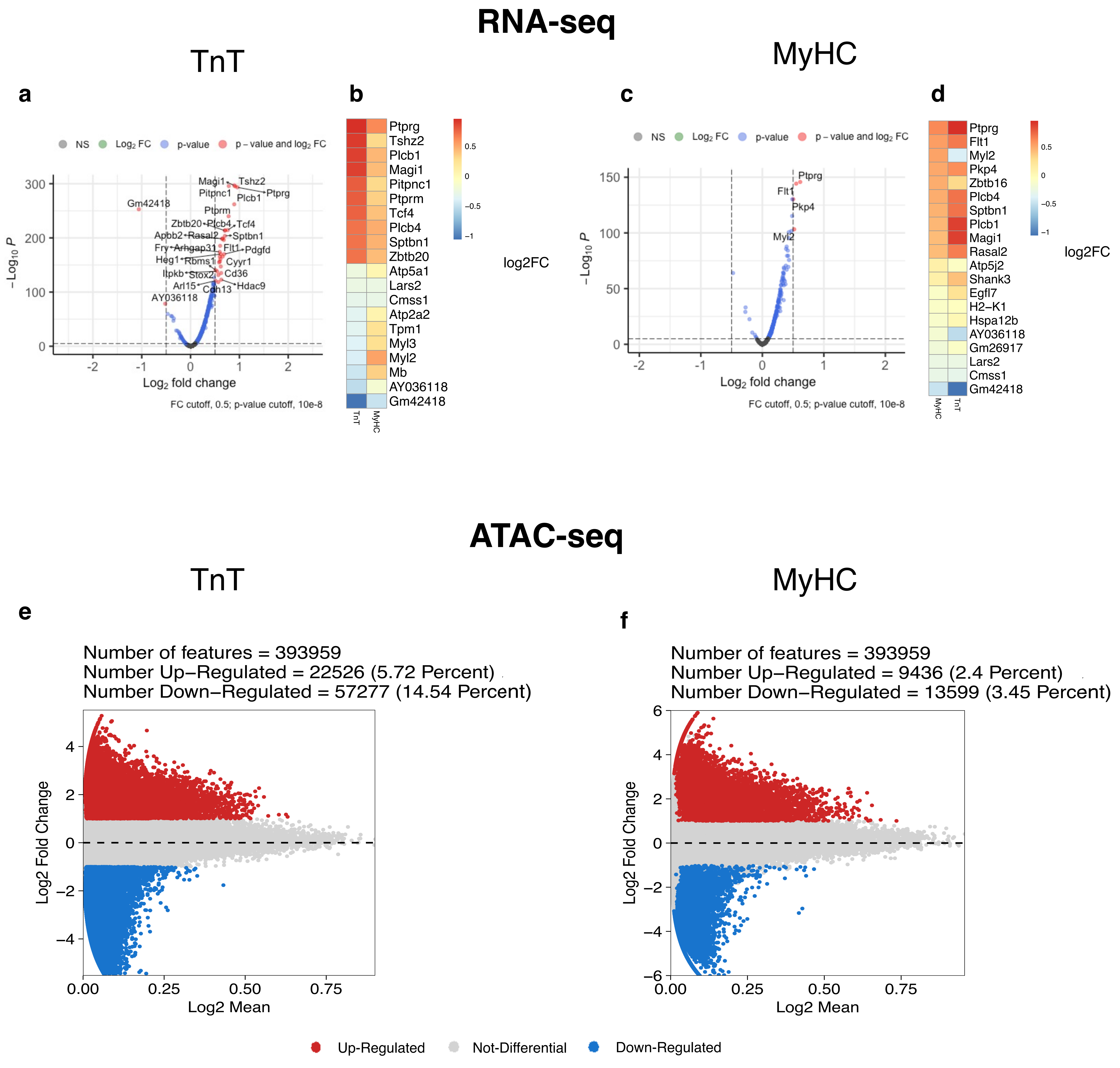


**Supplemental Figure 5**. **Gene expression and chromatin accessibility analysis of endothelial cell cluster C14.**

**a-d.** RNA-seq: Volcano plots and heat maps of DEGs (adj p<0.05) for mutant TnT and MyHC ECs compared to controls **e-f.** ATAC-seq: Differentially-accessible features in mutant TnT and MyHC ECs compared to controls (FDR ≤ 0.05, Log2FC ≥ 1).

**Suppl. Fig 6**

#### C14-Endothelial Cells

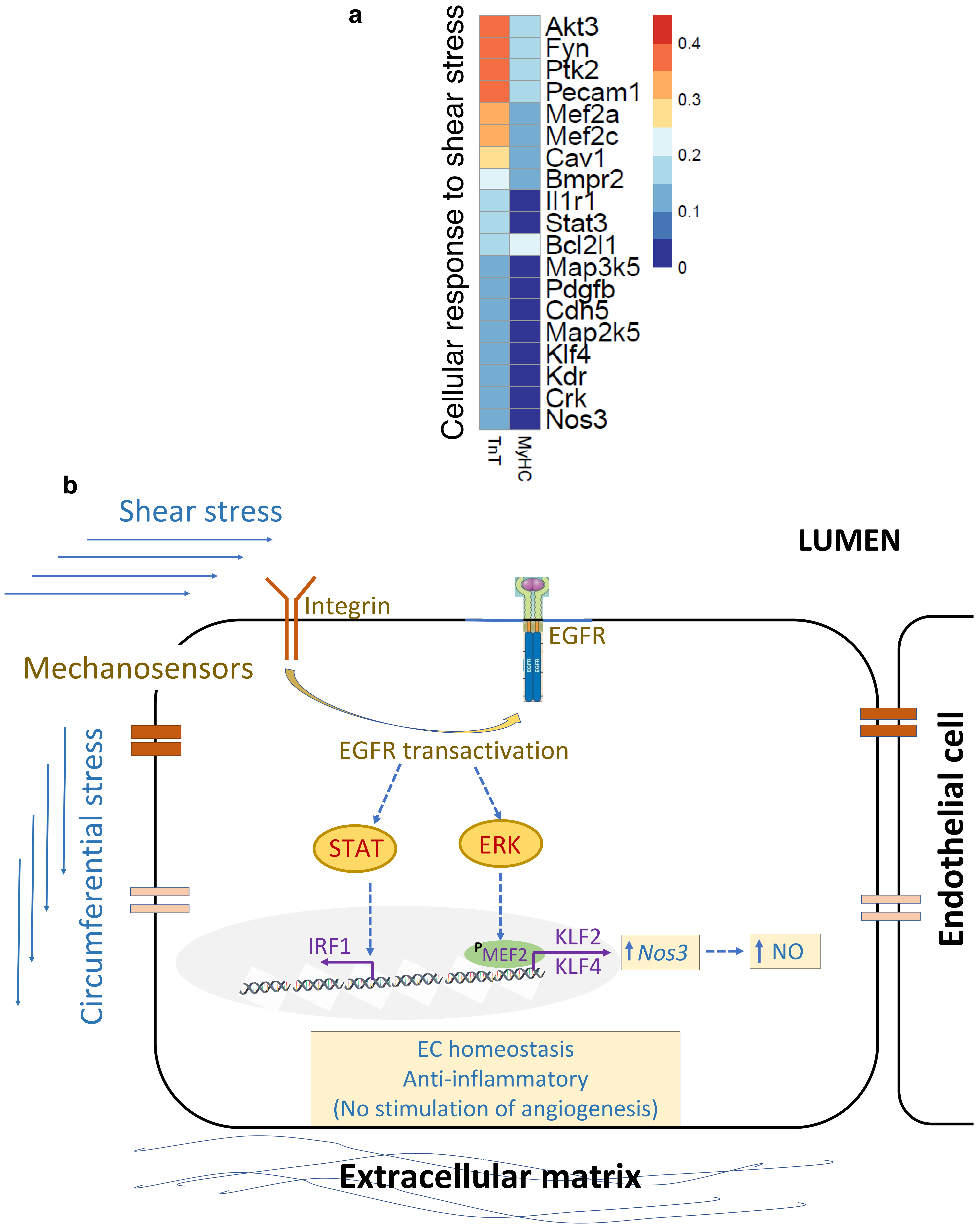


**Supplemental Figure 6**

**a.** Analysis of genes involved in cellular response to shear stress (AmiGO gene list) reveal greater dysregulation of gene expression in mutant TnT when compared to mutant MyHC ECs. **b. Schematic.** Our analysis of RNA-seq and ATAC-seq data (**Figs 4c,d, g-k, Suppl Fig 6a**) leads us to hypothesize that hyperdynamic LV function increases shear stress and circumferential stress in coronaries, which activates mechanosensitive receptors such as integrins, FAK signaling which leads to EGFR transactivation. This leads to activation of STAT and ERK signaling, that promote Mef2 phosphorylation and transcription of Klf2/4, which leads to activation of downstream genes such as NOS3 and increased NO production by ECs.

**Suppl. Fig 7**

### C2 - Macrophages

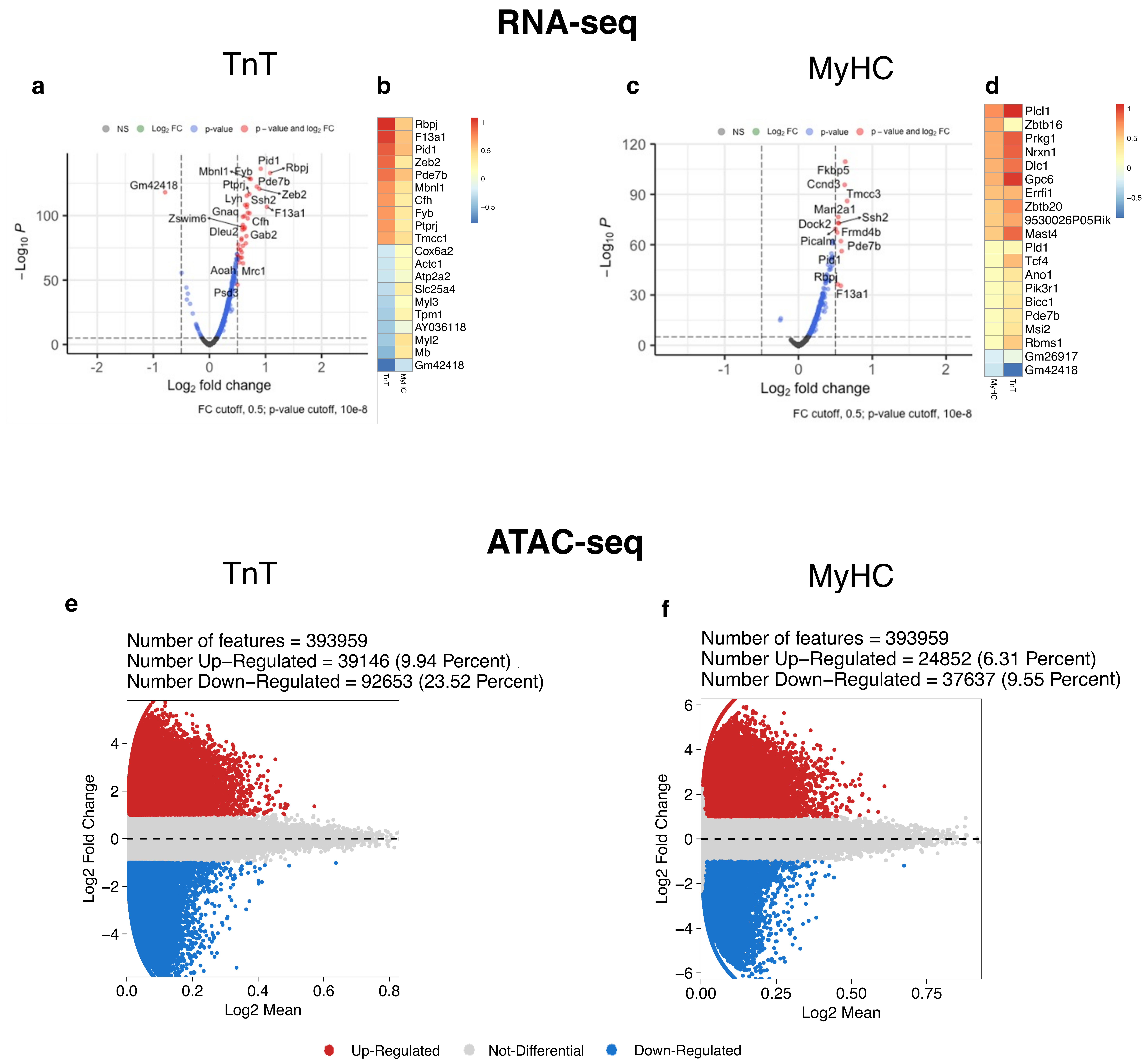


**Supplemental Figure 7. Cardiac macrophages (C2).**

**a-d.** RNA-seq: Volcano plots and heat maps of DEGs (adj p<0.05) in mutants compared to controls. **e, f.** ATAC-seq: Differentially-accessible features ((FDR ≤ 0.05, log2FC ≥ 1) in mutant TnT and MyHC show greater changes in mutant TnT.

**Suppl. Fig 8** 

### C2 - Macrophages

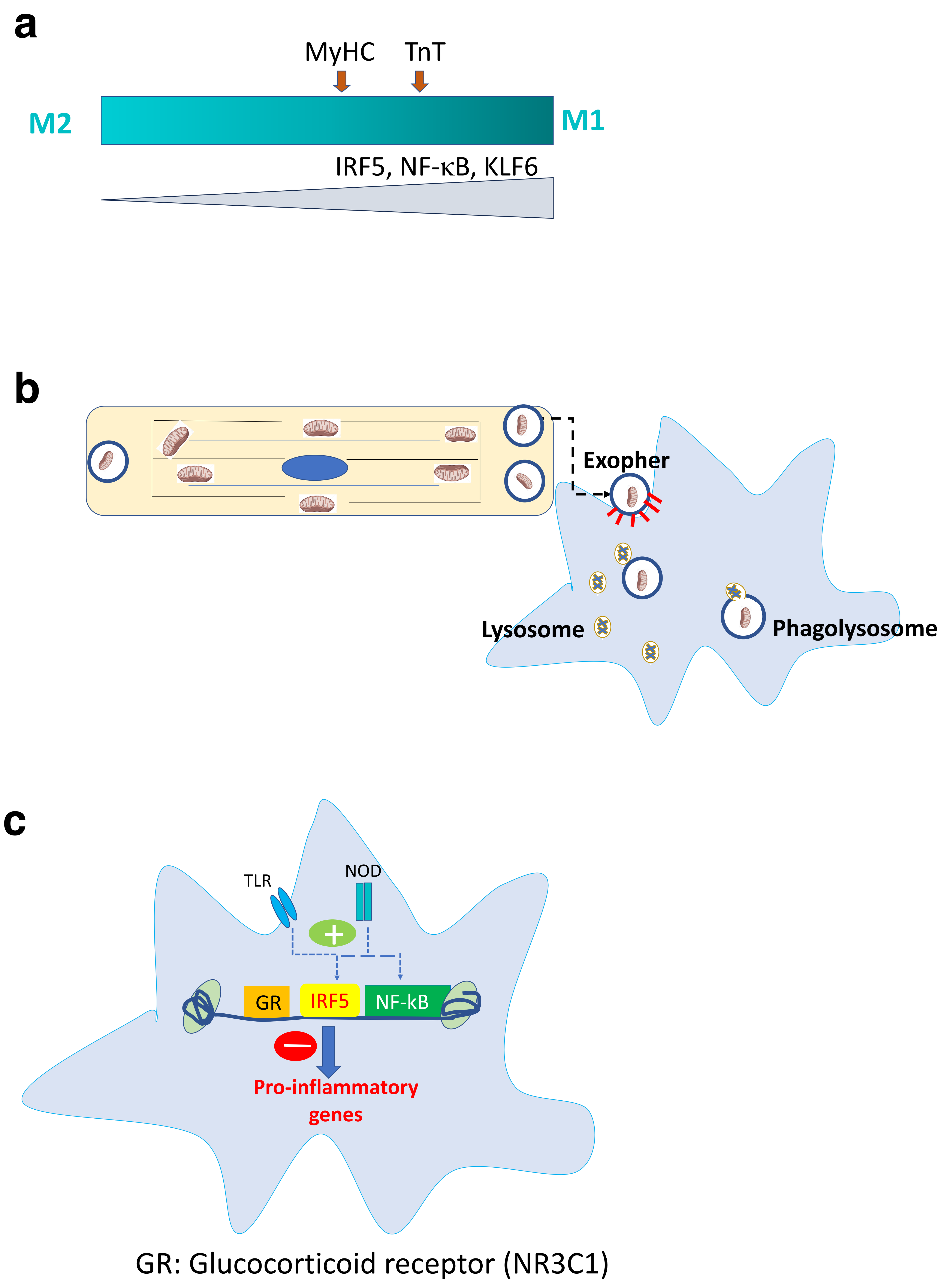


**Supplemental Figure 8. Macrophage Schematic.**

Our analysis of RNA- and ATAC-seq data in cardiac macrophages leads us to hypothesize the following. **a.** Greater activation of IRF5, REL (NFkB), IRF regulons in mutant TnT macrophages (**Fig 6h**) suggest greater macrophage polarization[101] towards M1 in mutant TnT when compared to mutant MyHC. **b.** Damaged mitochondria and other cargo from mutant myocytes is phagocytosed by resident cardiac macrophages and degraded by phagolysosomes (**Fig 6c-d**, **k**), which promotes cardiac myocyte homeostasis[59]. **c.** Activation of pattern recognition receptors such as TLR (toll-like receptors) and NOD (nucleotide binding and oligomerization domain-like receptors) by damaged mitochondria from cardiac myocytes, leads to nuclear translocation and DNA binding of IRFs, NF-kB (REL) (**Fig 6h-i**) which is pro-inflammatory. Simultaneous activation of the glucocorticoid receptor (NR3C1) in mutant macrophages (**Fig 6h**) blocks the pro-inflammatory effects of IRF and NF-kB activation.

**Suppl. Fig 9**

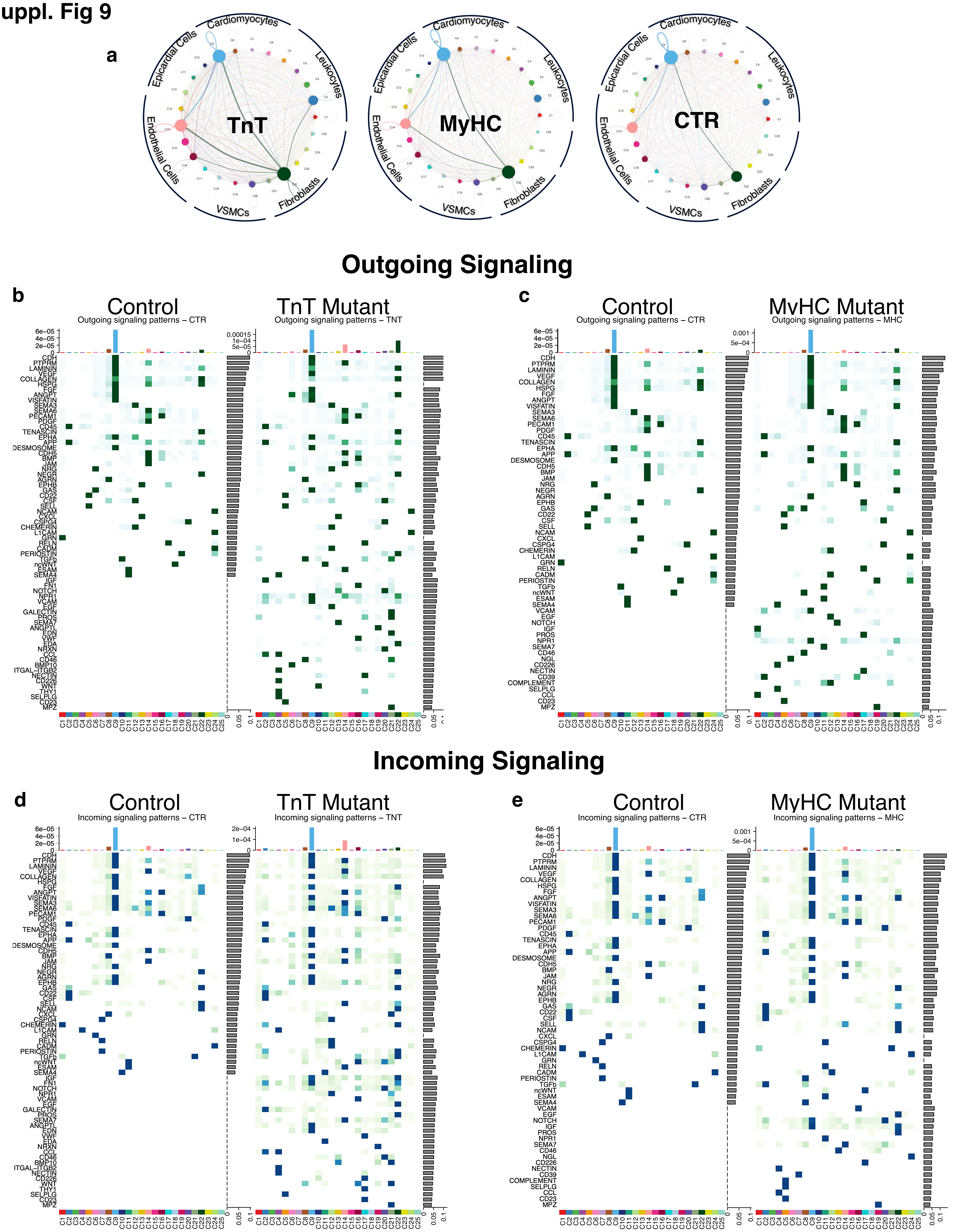


**Supplemental Figure 9. Cell-Cell-Communication Analysis with CellChat (RNA-seq).**

**a.** Communication between clusters shown for both mutants and the combined controls. Colored dots represent each cluster with color of connecting line showing origin of signaling. Cell types are grouped together. Stronger and added connections in both mutants compared to controls. **b, c.** Heatmap showing predicted outgoing signaling in controls and mutants for each detected signaling pathway. The left lower quadrant shows signaling that is only activated in the mutants can be seen. **d, e.** Heatmap showing predicted incoming signaling in controls and mutants for each detected signaling pathway. The left lower quadrant shows signaling that is only activated in the mutants can be seen.

### Suppl. Fig 10

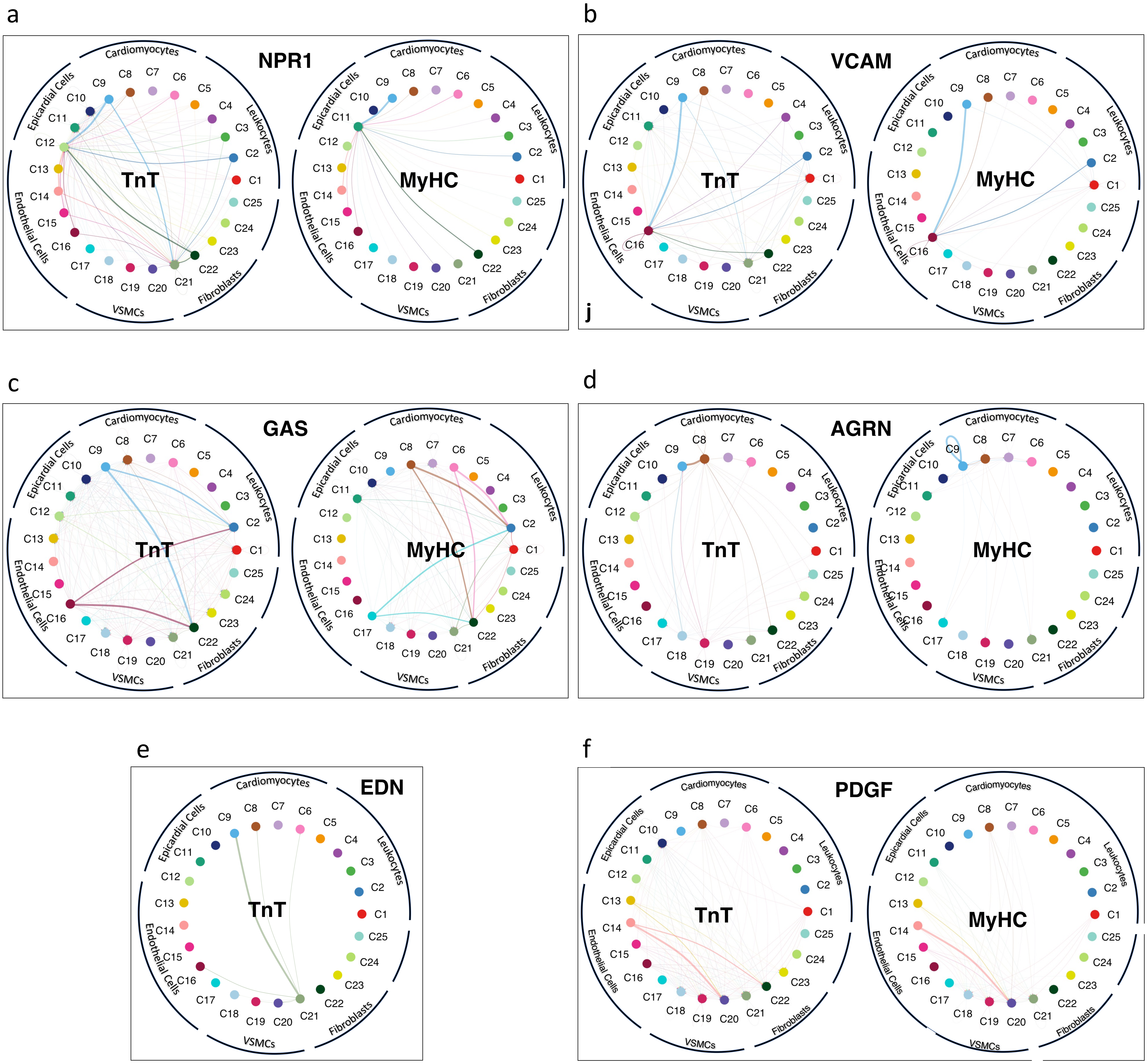


**Supplemental Figure 10. RNA-seq: Comparison of predicted signaling in mutants for NPR1, VCAM, GAS, AGRN, EDN, PDGF. Clusters are represented as dots and line color depicts the originating cluster of the depicted signaling.**

**a.** NPR1 signaling is predicted to originate from several clusters, with C9 and C22 being the major senders; signaling is received by the epicardial cell cluster C12 and the fibroblast cluster C21 in mutant TnT, and by the epicardial cell cluster C11 in mutant MyHC. **b.** VCAM signaling is predicted to originate from several clusters, and is received by C16 in both mutants. **c.** GAS signaling is predicted to originate from C9 and C16 in the TnT mutant, and from C8 and C17 in the MyHC mutant; major signaling recipients are the macrophage cluster C2 and the fibroblast cluster C22, in both mutants. **d.** AGRN signaling originates from C8 in mutant TnT, and C9 in mutant MyHC; main recipient in mutant TnT is the C8 cardiomyocyte cluster and C9 (autocrine signaling) in mutant MyHC. **e.** EDN (endothelin) signaling is only predicted in TnT. EDN signaling originates from the fibroblast cluster C21, and is received by the ventricular cardiomyocyte cluster C9. **f.** PDGF signaling mainly originates from endothelial cell cluster C14 and is received by the VSMC cell cluster C20 in both mutants, and the fibroblast cluster C22 only in TnT.

**Suppl. Fig 11**

### C22 - Fibroblasts

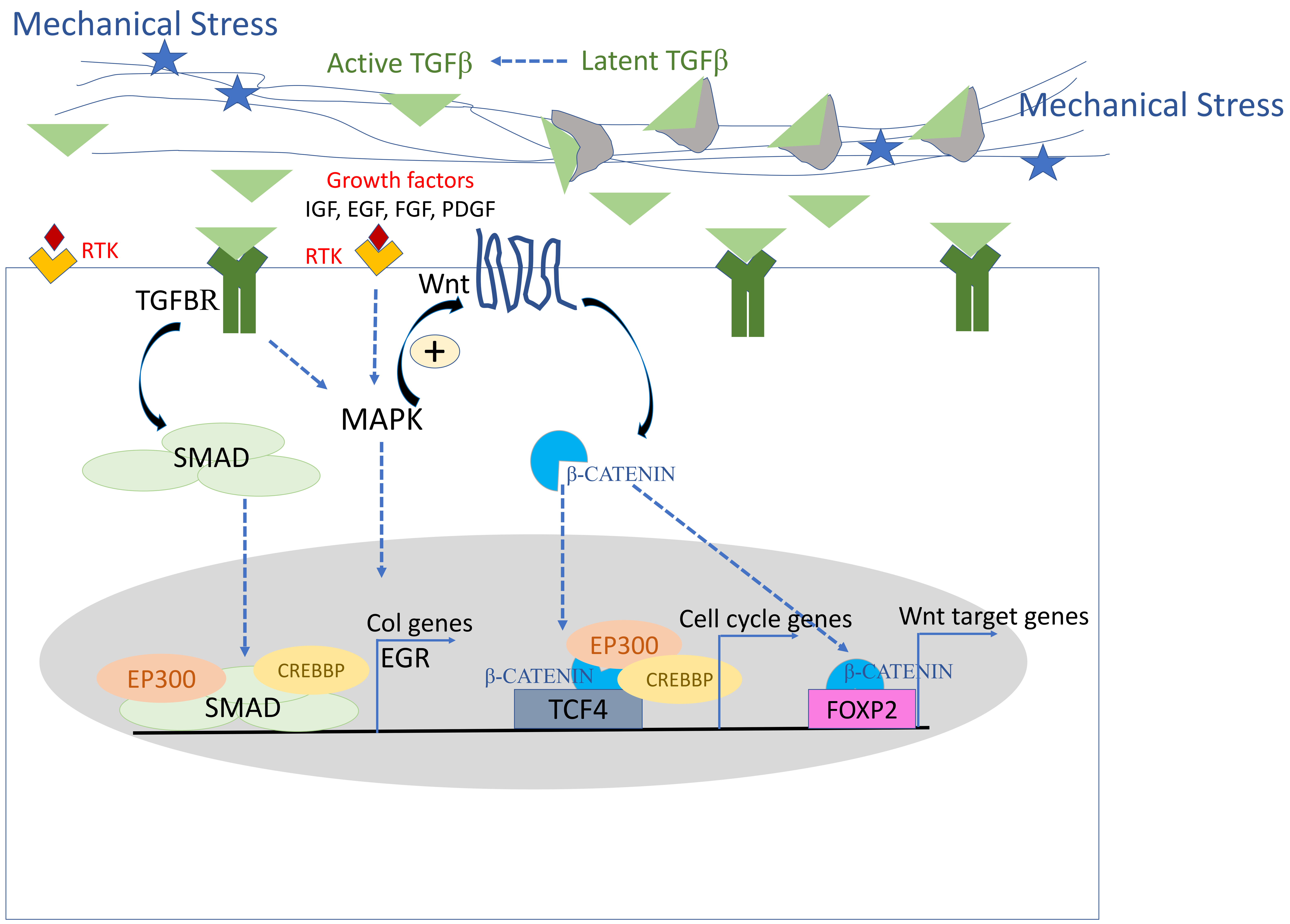


**Supplemental Figure 11**. **Schematic for myofibroblast activation and stimulation of interstitial fibrosis in mutant TnT hearts.** Our RNA-/ATAC-seq and echocardiography data led us to hypothesize that hyperdynamic LV function/diastolic dysfunction (**Fig 1d**) promote release of active TGFß in from cardiac extracellular matrix. TGFß binds TGFBR on fibroblasts, which activates SMADs (**Fig 3d**) and MAPK signaling (**Fig 3**j). Activated SMADs translocate to the nucleus where they associate with cofactors such as EP300, CREBBP to induce expression of collagen genes[87] **(Fig 3d**). Growth factors such as IGF, EGF, FGF, PDGF can also activate MAPK signaling (**Fig 3j**) by binding receptor tyrosine kinase (RTK) receptors on fibroblasts. MAPK activation (**Fig3d**) can stimulate Wnt-ß-catenin signaling. Beta-catenin translocates to the nucleus, where it associates with TCF4 and FOXP2 (**Fig 3d-i**) to promote transcription of cell cycle genes and Wnt target genes[88], thus potentiating the pro-fibrotic effect of activated TGFβ signaling.

#### Suppl. Fig 12

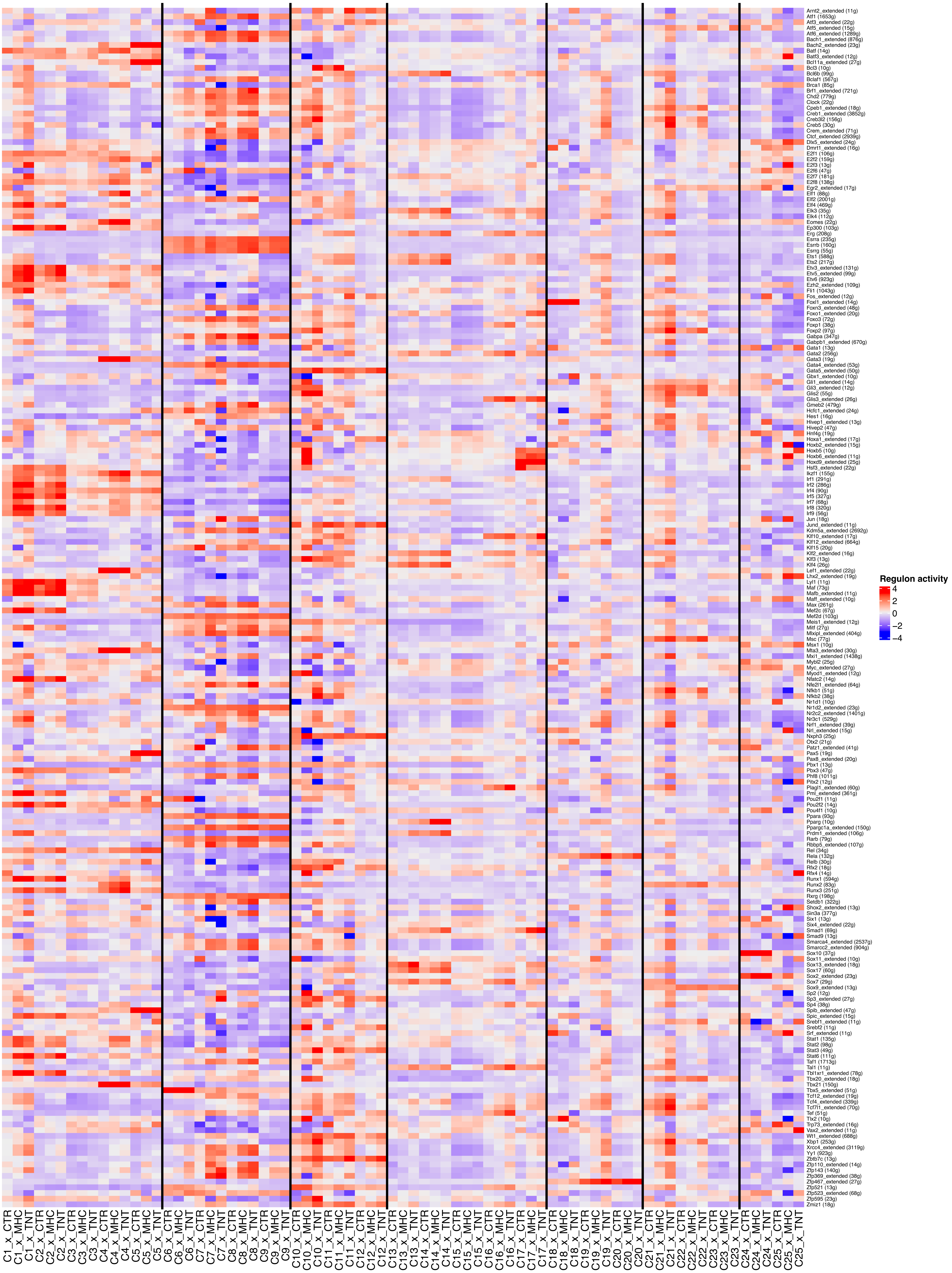

**Supplemental Figure 12**.

Regulon activity across all clusters and experimental conditions. Names of regulons are given on the right with the number of regulated genes in parenthesis. Genes are shown in alphabetical order. Individual cell types are delineated with black lines.
